# IFNA pathway drives the more aggressive phenotype of *KRAS*^G12D^-mutant pancreatic ductal adenocarcinomas via IFNAR1/STAT3 activation

**DOI:** 10.1101/2022.06.29.497540

**Authors:** Koetsu Inoue, Daniel H. Schanne, Aya Matsui, Pinji Lei, Sebastian Klein, Shuichi Aoki, Hajime Taniguchi, Hiroto Kikuchi, Jiang Chen, Zelong Liu, Shengdar Q. Tsai, Tyge CE Schmidt, Masaaki Iwasaki, Glenn Geidel, Alexander Koch, Peigen Huang, Dai Fukumura, Toshihiro Shioda, Lance L. Munn, Carlos Fernandez-del Castillo, Theodore S. Hong, Rakesh K. Jain, Andrew Liss, Nabeel Bardeesy, Dan G. Duda

## Abstract

Activating mutations of *KRAS* play critical roles in the initiation and progression of pancreatic ductal adenocarcinoma (PDAC). Accumulating evidence indicates that distinct *KRAS* alleles associate with different prognoses, but the underlying mechanisms are not known. We established isogenic *KRAS* mutants (*KRAS*^G12D^, *KRAS*^G12V^, and *KRAS*^WT^) using a *KRAS*^G12R^ patient-derived PDAC cell line by CRISPR/Cas9 knock-in. We used these isogenic cell lines, a collection of characterized human PDAC patient-derived cell lines, and murine PDAC models to study the role of these *KRAS* alleles *in vitro* and *in vivo*. We verified that the growth of *KRAS*^G12D^ cells is more aggressive compared to *KRAS*^G12V^ isogenic cells *in vitro* and *in vivo* using orthotopic mouse models. Signal transducer and activator of transcription (STAT) activation was the most significant difference between *KRAS*^G12D^ and *KRAS*^G12V^ isogenic PDACs. Furthermore, activation of interferon-alpha (IFNA)/IFNA receptor (IFNAR)1/STAT3 signaling in the cancer cells mediated the more aggressive phenotype of *KRAS*^G12D^ PDACs. Conversely, inhibition of IFNAR1 in patient-derived PDAC cells suppressed tumor growth. Finally, IFNAR1 blockade was also effective in murine PDAC models and induced a significant increase in survival when combined with immune checkpoint blockade therapy. We conclude that the IFNA pathway and IFNAR1/STAT3 axis contribute to a more aggressive tumor progression in human *KRAS*^G12D^ PDACs and that IFNAR1 inhibition is a potential therapeutic target for overcoming resistance to immunotherapy in PDAC.

**One Sentence Summary:** IFNA pathway drives the more aggressive phenotype of *KRAS*^G12D^-mutant pancreatic ductal adenocarcinomas via IFNAR1/STAT3 activation.

## INTRODUCTION

Pancreatic malignancies are the fourth most common cause of cancer-related death in the United States, with an increased incidence and continued unfavorable prognosis. The most aggressive and prevalent subtype—pancreatic ductal adenocarcinoma (PDAC)—has a 5-year overall survival rates of approximately 10% (*1, 2*). PDAC will become the second-leading cause of cancer-related death by 2030 (*3*). Significant clinical and preclinical research efforts over the last decades have resulted in a limited increase in long-term survival in PDAC patients so far (*4*).

One of the defining biological features of PDAC is an activating mutation of *KRAS*. More than 90% of PDACs carry a point mutation in codon 12 that leads to a switch of amino acids from glycine to aspartate (G12D, 51%), valine (G12V, 30%), or arginine (G12R, 12%) (*5, 6*). These events cause *KRAS* to be in a constitutively active state, which steers the affected cells towards a malignant phenotype (*7*). In PDAC, *KRAS* is one of the principal drivers of the disease through its involvement in signaling pathways that promote migration, cell proliferation, metabolism, and interaction with the tumor microenvironment (*8–11*). This understanding has led to major efforts to develop *KRAS* inhibitors. Recently, *KRAS*^G12C^ inhibitors have shown promising anti-tumor efficacy (*12, 13*). However, only 1% of PDACs carry the *KRAS*^G12C^ mutation (*14*). Effectively targeting more frequent mutations, such as *KRAS*^G12D^, remains an unmet need.

Several clinical studies have investigated whether different *KRAS* mutations are associated with distinct clinical PDAC outcomes (*15*). In a recently reported phase 1/2 study of neoadjuvant radio-chemotherapy in 50 patients with resectable PDAC, we found a statistically significant lower overall survival (OS) in patients with *KRAS*^G12D^ tumors compared to other mutations or wild-type *KRAS* (*16*). In another study, Ogura *et al*. screened a group of 242 biopsies from unresectable PDACs patients. They found that the patients with *KRAS*^G12D^ PDACs and *KRAS*^G12R^ mutations had a worse prognosis (*17*). Finally, a more recent study of 219 European patients with advanced PDAC showed the same association between *KRAS*^G12D^ mutation and shorter OS (*18*). Thus, characterizing the biological consequences of different *KRAS* mutations could provide new insights into tumor pathophysiology and reveal specific vulnerabilities in PDAC and in other tumors with frequent *KRAS* mutations, such as colon and non-small cell lung cancer. Based on these data, we hypothesized that the type of *KRAS* mutation differentially mediates tumor progression and treatment resistance.

This hypothesis could not be directly tested previously. Previous preclinical studies used xenograft-derived cell lines or animal models carrying a *KRAS*^G12D^ mutation. However, these cell lines may carry multiple genetic or epigenetic alterations, making it difficult to precisely identify how various biological features associate with distinct *KRAS* mutations. To address this limitation, we generated isogenic PDAC patient-derived cells via CRISPR/Cas9 knock-in. These well-defined, genetically engineered models served as a platform to study the role of different *KRAS* mutations, along with a panel of other patient-derived PDAC cell lines and murine models. Using these models, we evaluated PDAC growth *in vitro* and *in vivo*, examined the causality between changes in downstream targets, and studied the impact of targeting them, genetically and pharmacologically, in orthotopic models of PDAC in mice.

## RESULTS

### Isogenic cell lines with different *KRAS* alleles show differential growth rates *in vitro* and *in vivo*

To determine whether the *KRAS* allele type mediates PDAC progression, we first generated isogenic cell lines from the PDX-derived cell line (PDCL)-1108, which harbored a *KRAS*^G12R^ mutation. Briefly, we introduced Cas9 protein along with sgRNA’s targeting *KRAS* exon 2 and single-stranded DNA donor templates coding for alternative *KRAS* alleles (*KRAS*^WT^, *KRAS*^G12D^, and *KRAS*^G12V^); we then analyzed single-cell clones for successful integration by restriction digest, Sanger sequencing and deep sequencing (**Figs. 1A** & **S1**).

**Fig. 1:**
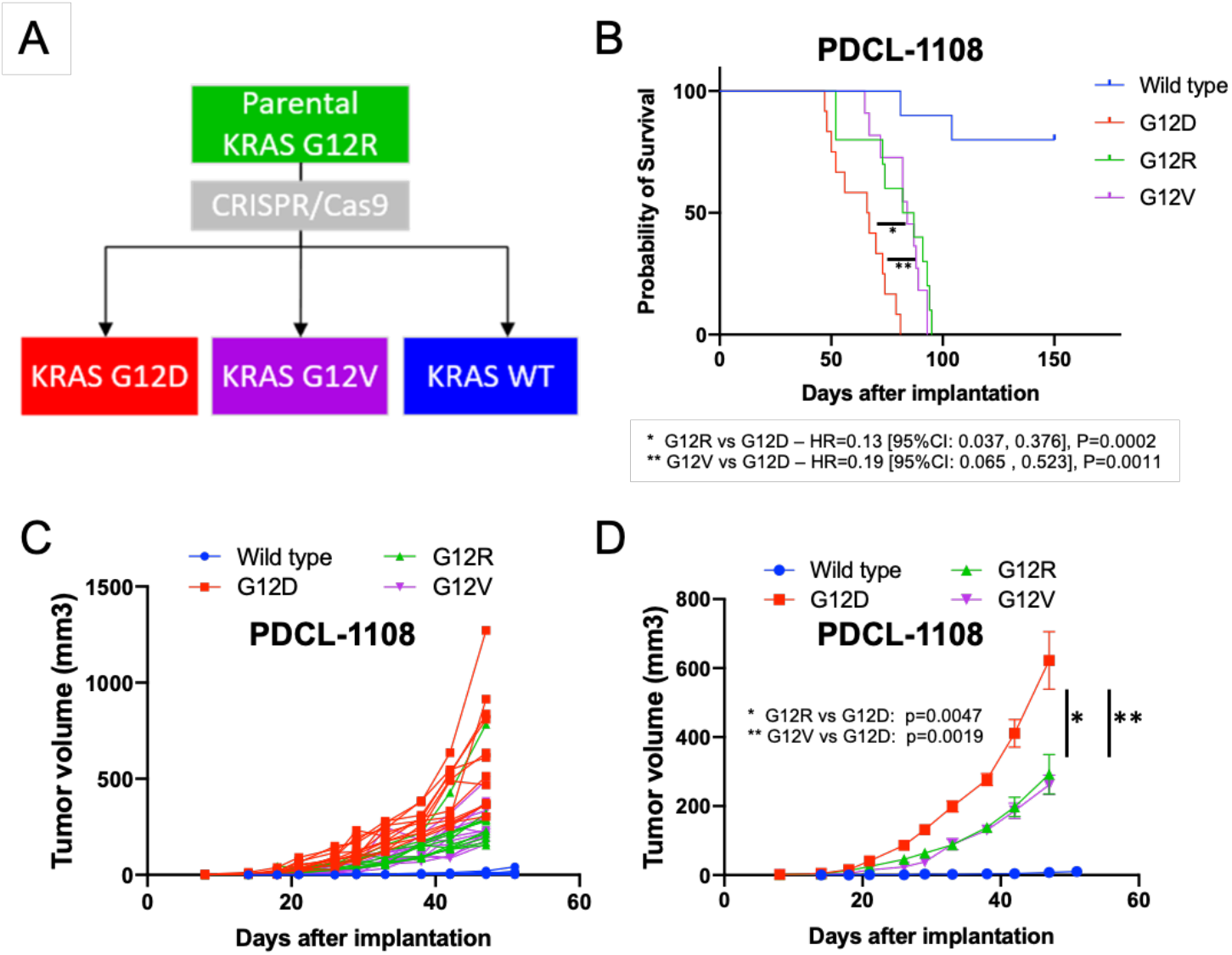
Isogenic PDAC cells with different *KRAS* alleles show differential tumor progression. (**A**) Schema showing isogenic cell lines established by CRISPR/Cas9 technology. (**B**) Kaplan-Meier curve depicting mouse survival. Individual growth curves in (**C**) and average tumor size with SEM in surviving mice in (**D**). Cancer cells were implanted into NSG-human-HGF-knock-in mice. G12D: n=12; G12R: n=10; G12V: n=12; wild type: n=10; n refers to biological replicates. Tukey’s test for tumor volume and Cox regression test for Kaplan-Meier survival distributions.

To assess whether the isogenic cell lines exhibit differential growth rates *in vitro* and *in vivo*, we performed 2-*D* and 3-*D* proliferation assays and evaluated tumor progression in orthotopic PDAC mouse models. *KRAS*^G12D^ cells showed enhanced proliferation capacity compared to *KRAS*^G12V^ and *KRAS*^WT^ *in vitro* (**Fig. S2**). Moreover, orthotopically grafted *KRAS*^G12D^ cells showed an accelerated tumor growth rate and the mice had shorter survival than those implanted with other mutants or wild-type tumors *in vivo* (**Fig. 1B-D**). The difference was most significant and reproducible when comparing *KRAS*^G12D^ and *KRAS*^G12V^ PDAC cells in the orthotopic model in mice. Thus, we focused our further studies on these two *KRAS* alleles.

### Isogenic cell lines with different *KRAS* alleles show similar levels of expression of *KRAS* and downstream targets, and comparable sensitivity to MAPK or PI3K inhibition

Because previous reports showed that *KRAS* copy number gain is associated with outcome in human PDAC (*7, 19*), we next checked *KRAS* expression levels between the isogenic cell lines. We found no significant differences (**Fig. S3A**). We further assessed whether different *KRAS* allele types have differential phosphorylation of canonical downstream targets such as ERK or AKT by Western blotting (*20, 21*). We found no significant differences between the isogenic cell lines (**Fig. S3B-D**). In addition, it has been reported that different *KRAS* alleles may have differential sensitivity to MEK inhibitors (*19*). Therefore, we checked drug sensitivity to BRAF, MEK, ERK, and PI3K inhibitors. We found no significant differences in drug sensitivity among isogenic cell lines (**Fig. S3E-H**). These results indicate that non-canonical mechanisms may contribute to the differential tumor progression between isogenic PDCLs with different *KRAS* alleles.

### STAT3 is activated, and STAT1 is suppressed in *KRAS*^G12D^ PDACs

To reveal differentially activated pathways in PDCLs with distinct *KRAS* alleles *in vivo*, we performed bulk-tissue RNA sequencing (RNA-seq). Analysis was performed using size-matched *KRAS*^G12D^ (n=3) and *KRAS*^G12V^ (n=3) tumor samples, all collected when the tumors reached 8mm in diameter. Gene Set Enrichment Analysis (GSEA) showed that *“regulation of peptidyl serine phosphorylation of STAT protein”* was the most significantly enriched gene set in the *KRAS*^G12D^ compared to *KRAS*^G12V^ tumors (**Fig. 2A, B**). The heatmap of the *“regulation of peptidyl serine phosphorylation of STAT protein”* gene set indicated upregulated expression levels of *IFNA1* and *IFNA13* in the *KRAS*^G12D^ tumors (**Fig. 2C**). We validated that *KRAS*^G12D^ PDCLs had higher *IFNA13* expression *in vitro* and *in vivo* by real-time qPCR (**Fig. 2D**).

**Fig. 2:**
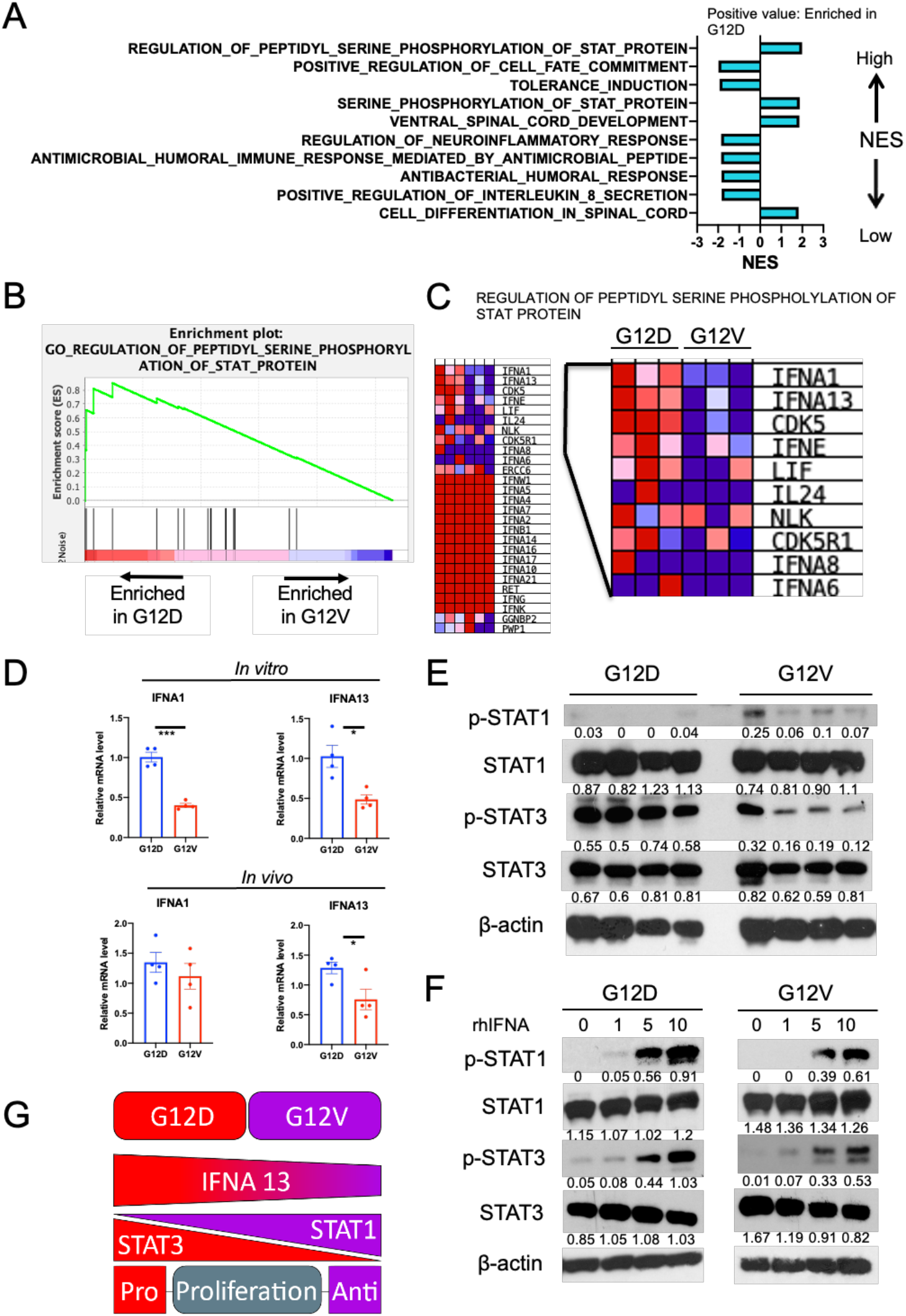
Interferon alpha (IFNA) mediates the activation of STAT3 in *KRAS*^G12D^ PDXs of PDAC. (**A**-**C**) Gene Set Enrichment Analysis (GSEA) analysis of RNA-seq data (G12D versus G12V); data mapped to the human genome (hg38). G12D: n=3; G12V: n=3; n refers to biological replicates. (**A**) Gene Ontology (GO) term list showing top ten GO terms according to Normalized Enrichment Score (NES). (**B**) Enrichment plot of regulation of peptidyl serine phosphorylation of STAT protein. NES: 1.99, False discovery rate (FDR): 0.07 (**C**) Heatmap of “regulation of peptidyl serine phosphorylation of STAT protein” gene set enriched in *KRAS*^G12D^. Red, high, blue, low. (**D**) Human IFNA 1 and IFNA13 expression in cells and tumor tissues measured by real-time qPCR. Mean relative mRNA level is indicated with error bars representing SEM. All *in vitro* assays, n=3-4; for *in vivo* analyses n=4. *p<0.05, ***p<0.001 from Student’s t test. (**E**) STATs expression in tumor tissue. Total and phosphorylated STAT1 and STAT3 were measured by Western blotting. G12D: n=4; G12V: n=4. Representative of two or more independent experiments. (**F**) Response of STATs to different doses of recombinant human IFNA. Total and phosphorylated STAT1and STAT3 in cells were measured by Western blotting. Cells were treated with different concentration of recombinant human IFNA. Representative of two or more independent experiments. (**G**) Schematic representation showing STAT1, STAT3 and IFNA expression between the G12D and G12V alleles.

We next examined STAT1 and STAT3 activation, which are activated downstream IFNA (*22–24*).Western blotting analysis showed the increased STAT3 activation in the *KRAS*^G12D^ versus *KRAS*^G12V^ PDX PDAC tissues (**Fig. 2E**). Moreover, STAT3 activation was verified using a panel of *KRAS*^G12D^ versus *KRAS*^G12V^ PDX PDAC cells (**Fig. S4**). Exposure to exogenous recombinant human (rh)IFNA confirmed activation of both STAT1 and STAT3 at high concentrations (**Fig. 2F**).

STAT1 and STAT3 activation could be reciprocally regulated and have opposing effects on tumor progression (*25, 26*). Our results showed that increased IFNA expression in *KRAS*^G12D^ PDAC cells associated with more rapid tumor progression, activation of STAT3, and suppression of STAT1 activation. To further confirm these findings, we conducted a gene ontology (GO) annotation of the significantly differentially expressed genes between *KRAS*^G12D^ and *KRAS*^G12V^ PDACs (see details in **Methods**). We found that GO terms related to STAT1 signaling, including *response to virus, defense response to virus*, *type I IFN signaling pathway*, and *cellular response to IFNA*, were all enriched in *KRAS*^G12V^ PDACs (**Fig. S5A**). We also validated the results of the RNA-seq analysis for genes involved in *“response to virus”*, including IFIT2 and IFIT3, utilizing real-time PCR (**Fig. S5B**-**C**). In addition, we found that TRAIL, a downstream gene for STAT1 signaling, was among the top 5 differentially expressed genes (DEG) (**Fig. S5D**) and further verified its increased expression in *KRAS*^G12V^ tumors by real-time qPCR (**Fig. S5E**). These results show activation of STAT1 signaling in *KRAS*^G12V^ compared to *KRAS*^G12D^PDACs.

The STAT3 pathway leads to downstream NF-κB activation (*27*). We found that NF-κB activation in *KRAS*^G12D^ PDAC tissues by Western blotting (**Fig. S5F**). Taken together, our studies of isogenic PDCLs show that STAT3 is activated and STAT1 is suppressed in the more aggressive *KRAS*^G12D^ PDACs relative to the more indolent *KRAS*^G12V^ isogenic tumors (**Fig. 2G**).

### Genetic inhibition of IFNAR1 delays tumor progression in both *KRAS*^G12D^ and *KRAS*^G12V^ PDACs, and exogenous rhIFNA accelerates *KRAS*^G12V^ tumor growth

The IFNA/STAT1 axis may have inhibitory (*28, 29*) or promoting effects on tumor progression (*30, 31*). Thus, we next established IFNAR1-knock-down (KD) versions of the isogenic PDCLs to determine the role of the IFNA/STAT pathway in the PDAC models (**Fig. 3A**).

**Fig. 3:**
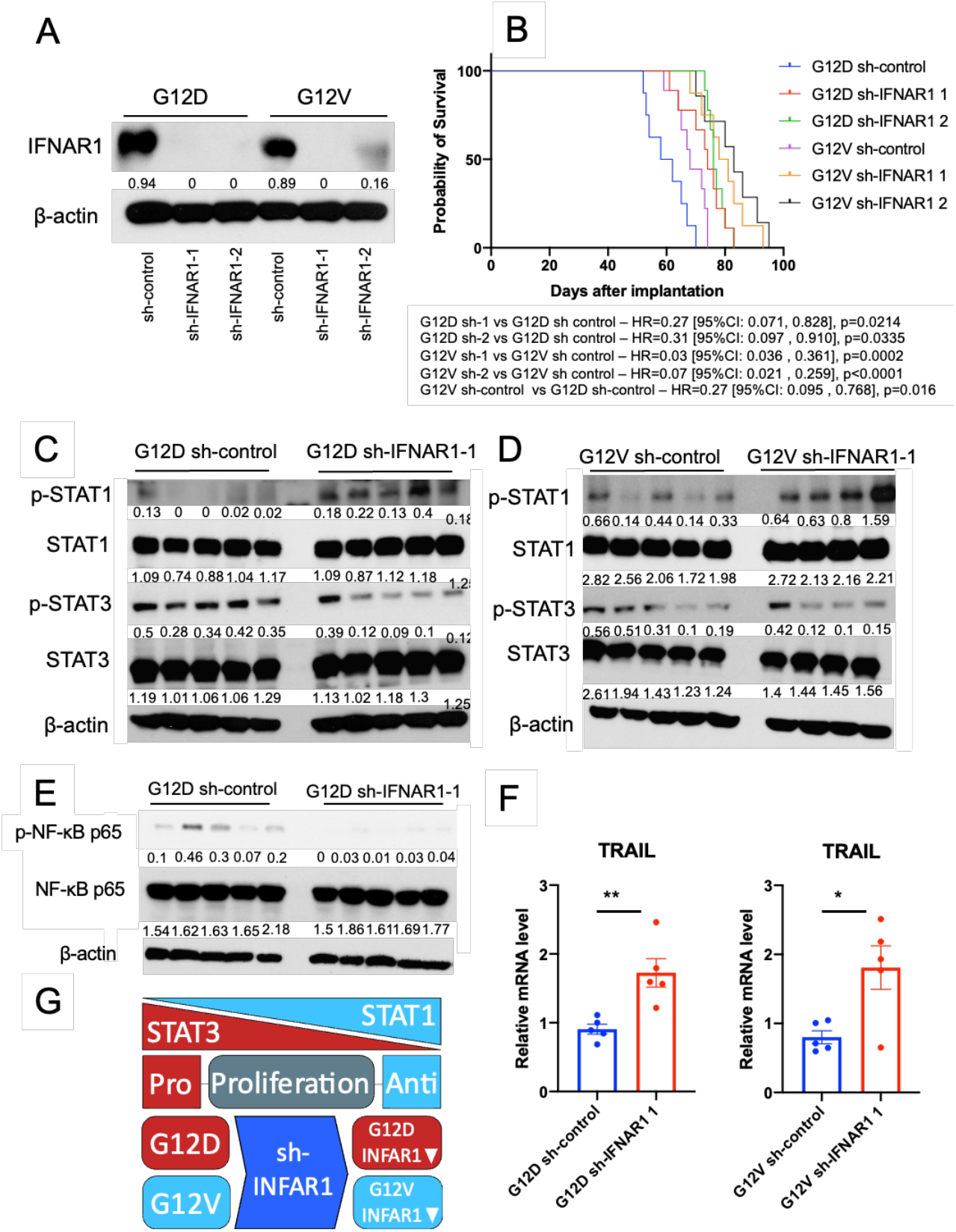
Genetic inhibition of interferon alpha receptor 1 (IFNAR1) regulates tumor progression *in vivo*. (**A**) Validation of IFNAR1 expression knockdown with Western blotting. Representative of two independent experiments. (**B**) Kaplan-Meier survival curves for orthotopic PDACs implanted in NSG-human-HGF-knock-in mice. G12D sh-control: n=8; G12D sh-IFNAR1 1: n=9; G12D sh-IFNAR1 2: n=9; G12V sh-control: n=9; G12V sh-IFNAR1 1: n=8; G12V sh-IFNAR1 2: n=7; n refers to biological replicates. p values from Cox regression test. (**C**) STATs expression in sh-control and sh-IFNAR1 1 tumor. Total and phosphorylated (p)-STAT1 and STAT3 were measured by western blotting. G12D sh-control: n=5; G12D sh-IFNAR1 1: n=5; G12V sh-control: n=5; G12V sh-IFNAR1 1: n=4. Representative of two or more independent experiments. (**D**) Total and p-NF-κB p65 expression in tumor. G12D sh-control: n=5; G12D sh-IFNAR1 1: n=5; G12V sh-control: n=5; G12V sh-IFNAR1 1: n=4. Representative of two or more independent experiments. (**E**) TRAIL expression in sh-control and sh-IFNAR1 1 tumor tissue measured by real-time qPCR. Mean relative mRNA level is indicated with error bars representing SEM. G12D sh-control: n=5; G12D sh-IFNAR1 1: n=5; G12V sh-control: n=5; G12V sh-IFNAR1 1: n=4. Assays were performed in triplicate or quadruplicate. *p<0.05, **p<0.01 from Student’s t test. (**F**) Schematic representation showing STATs status between sh-control and sh-IFNAR1.

We first confirmed that genetic IFNAR1 inhibition suppressed both the constitutive activation of STAT1 and STAT3 and that induced by rhIFNA, in both *KRAS*^G12D^ and *KRAS*^G12V^ IFNAR1-KD PDCLs (**Fig. S6A**). When we orthotopically implanted these PDCLs, we found that IFNAR1 inhibition delayed tumor growth and improved survival in both *KRAS*^G12D^ and *KRAS*^G12V^ IFNAR1-KD isogenic PDCL models (**Figs. 3B** and **S6B-C**).

Next, we conducted a separate time-matched study and assessed STAT activation in the tumor tissues. We found that IFNAR1 inhibition suppressed the activation of STAT3 and promoted STAT1 phosphorylation (**Figs. 3C-D**). In addition, genetic IFNAR1 inhibition suppressed NF-κB activation in *KRAS*^G12D^/IFNAR1-KD tumors (**Fig. 3E**). Although IFNA may activate ERK and AKT in cancer (*32*), IFNAR1 inhibition did not affect ERK and AKT activation in *KRAS*^G12D^/IFNAR1-KD PDACs (**Fig. S7A-B**). We further evaluated STAT1downstream genes by real-time qPCR. We found significant upregulation of TRAIL and other genes related to STAT1 signaling in IFNAR1-KD tumors, but not ERK (**Figs. 3F** and **S7C**). We also checked the effect of pharmacologic blockade with an anti-human IFNAR antibody *in vitro*. In line with data from the genetic approach, we found that blockade using an anti-human IFNAR antibody abolished STAT1 and STAT3 activation in both cell types (**Fig. S7D**). In addition, IFNAR antibody treatment reduced PDAC cell viability *in vitro* (**Fig. S7E**). These results show that IFNAR1 promotes tumor progression via STAT3/NF-κB activation and not via STAT1 activation in the *KRAS^G12D^* PDACs. Previous preclinical studies showed that rhIFNA treatment could suppress tumor growth (*33, 34*).However, when tested in clinical trials, combining rhIFNA with chemotherapy did not show benefits in PDAC patients (*35–38*). Thus, we treated mice bearing established orthotopic *KRAS*^G12D^ or *KRAS*^G12V^ PDAC tumors with rhIFNA or vehicle (sodium chloride solution) and examined the effect on mouse survival. Consistent with the clinical observations, we found no survival advantage in mice treated with rhIFNA, irrespective of *KRAS* allele type (**Fig. S8A,B**). Moreover, median OS in the rhIFNA-treated mice bearing *KRAS*^G12V^ PDAC tended to be shorter (by 10.5 days) compared to control-treated mice (85.5 days versus 96 days) (**Fig. S8B**) (p=0.061). These results show that IFNAR1 inhibition can delay tumor progression in both *KRAS*^G12D^ and *KRAS*^G12V^ models primarily via STAT3 suppression and suggest that reduced IFNA mediates the more indolent behavior of *KRAS*^G12V^ tumors compared to those that are more aggressive *KRAS*^G12D^ (**Fig. 3G**).

### STAT3 overexpression promotes tumor progression across *KRAS* mutated PDAC subtypes

We next tested the role of the IFNA/STAT3 axis in driving tumor progression. We first examined induced STAT3 overexpression in *KRAS*^G12D^ IFNAR1-KD cells (**Fig. 4A**). When we orthotopically implanted these tumor cells in mice, STAT3 overexpression reversed the inhibition of tumor growth (**Fig. 4B**).

**Fig. 4:**
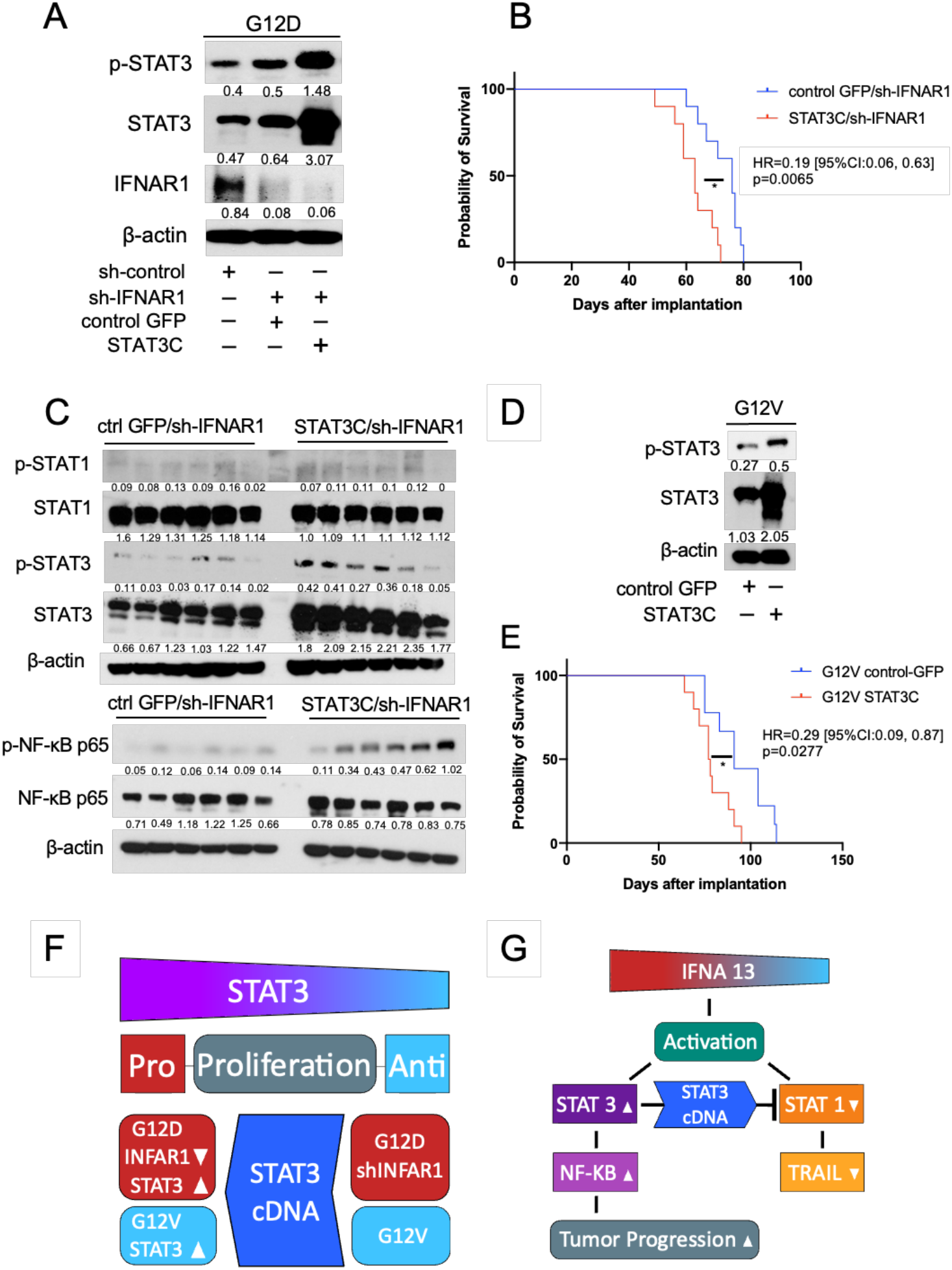
STAT3 overexpression promotes PDAC progression. **A**) Validation of STAT3 overexpression. IFNAR1-silenced *KRAS*^G12D^ cells were transfected with constitutively active STAT3 mutant (EF.STAT3C.Ubc.GFP) or control GFP (pLVE-eGFP). GFP positive cells were collected by cell sorting 7 days after transfection. Total and phosphorylated (P)-STAT3 and IFNAR1 in *KRAS*^G12D^ cells were measured by Western blotting. Representative of two or more independent experiments. (**B**) Kaplan-Meier survival distributions in NSG-human-HGF-knock-in mice bearing orthotopic PDAC. G12D control GFP/ sh-IFNAR1 1: n=10; G12D control GFP/ sh-IFNAR1 1: n=10; n refers to biological replicates. p values from Cox regression test. (**C**) Total and p-STAT1, STAT3 and NF-κB p65 expression level in tumor tissue. Control GFP/sh-IFNAR1: n=6; STAT3C/sh-IFNAR1: n=6. Representative of two or more independent experiments. (**D**) Validation of STAT3 overexpression. *KRAS*^G12V^ cells were transfected with constitutively active STAT3 mutant (EF.STAT3C.Ubc.GFP) or control GFP (pLVE-eGFP). Representative of two or more independent experiments. (**E**) survival distributions in NSG-human-HGF-knock-in mice bearing orthotopic PDAC. G12V control GFP: n=9; G12V STAT3C: n=10. p values from Cox regression test. (**F**) Schematic representation of STAT3 role in mutant *KRAS* subsets tumor progression. (**G**) Model indicating the mechanism by which *KRAS* alleles differentially mediate tumor progression.

Next, we evaluated total and phosphorylated STAT3 levels in separate time-matched studies and confirmed their overexpression in STAT3C/IFNAR1-KD tumors, and p-NF-κB upregulation (**Fig. 4C** and **S9**). To verify that the IFNA/STAT3 axis mediated the differential tumor growth rate between *KRAS*^G12D^ and *KRAS*^G12V^ PDCL, we examined whether STAT3 overexpression also rescues the delayed tumor progression in *KRAS*^G12V^ PDCL. Indeed, STAT3 overexpression in *KRAS*^G12V^ cells accelerated tumor progression in the orthotopic PDAC model (**Fig. 4D-E**).

We also tested whether STAT1 inhibition in IFNAR1-KD PDAC cells affects tumor growth by generating double knock-down lines for both IFNAR1 and STAT1 (**Fig. S10A**). We found no significant difference in tumor growth or mouse survival after orthotopic implantation of these PDAC cells (**Fig. S10B**). Consistent with these results, pharmacologic inhibition of STAT1 with fludarabine or the Janus kinase (JAK) 1/2 inhibitor ruxolitinib (Javaki) did not affect the viability of *KRAS*^G12D^ or *KRAS*^G12V^ cells *in vitro* (**Fig. S10C**). These results support the conclusion that the IFNAR1/STAT3 axis plays a critical role in PDAC and mediates the accelerated tumor growth in *KRAS*^G12D^ versus *KRAS*^G12V^ isogenic PDCL cells (**Fig. 4F-G**).

### IFNAR1 induces more aggressive growth in human *KRAS*^G12D^ PDAC models, and high levels associated with shorter survival in PDAC patients

To confirm that *KRAS*^G12D^ reproducibly shows higher IFNA expression level and STAT3 activation, we evaluated their expression across multiple PDX tumors. Analysis of RNA-seq data from 25 independent PDX tumors showed higher levels of IFNA in PDX tissues from PDAC with *KRAS*^G12D^ mutation, which were significant for *IFNA13* expression (**Fig. 5A**). In addition, an immunohistochemical evaluation showed higher levels of p-STAT3 in *KRAS*^G12D^ PDX tumor tissues (**Fig. 5B-C**). To evaluate whether IFNAR1 promotes tumor growth in these PDX models, we selected two PDACs based on differential IFNAR1 expression: PDCL-1319 (high levels) and (low levels) (**Fig. 5D**). PDCL-1319 and PDCL-609 *KRAS*^G12D^ cells had similar IFNA1 and IFNA13 expression levels (**Fig. S11A**). Moreover, we confirmed increased activation of STAT1 and STAT3 in response to IFNA in the cells with higher levels of IFNAR1 expression (PDCL-1319) (**Fig. 5E**). We next tested the effect of IFNAR1-KD in these PDCLs (**Fig. 5F**) to determine the impact of genetic IFNAR1 inhibition on STAT activation and tumor growth. Western blot analysis showed inhibition of STAT1 and STAT3 activation in PDCL-1319 IFNAR1-KD and PDCL-609 IFNAR1-KD cells (**Fig. S11B**). Furthermore, genetic IFNAR1 inhibition repressed tumor growth significantly in orthotopic PDAC mouse models (**Fig. 5G-H**). The inhibitory effects were more pronounced in the PDCL-1319 (high-IFNAR1) model.

**Fig. 5:**
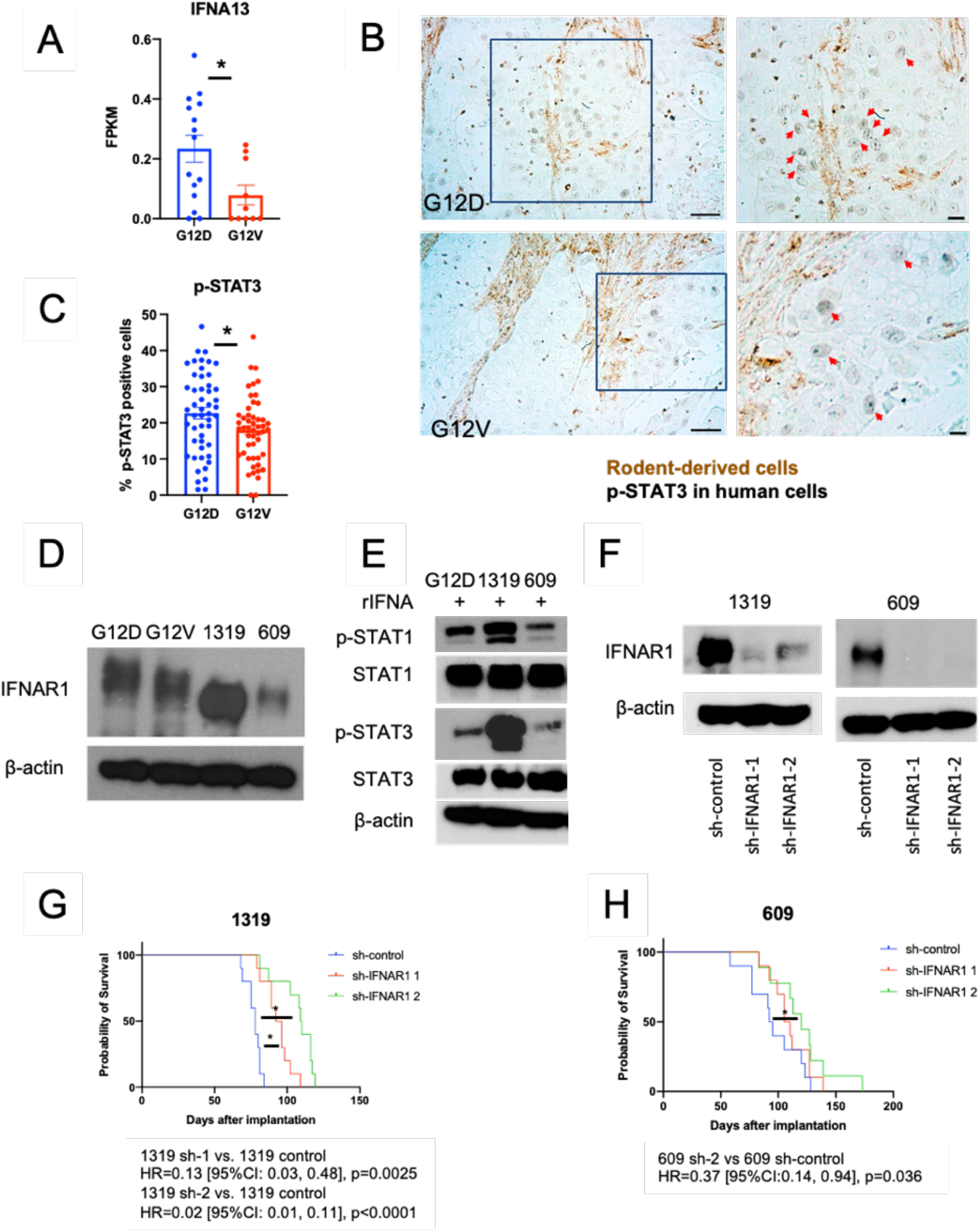
*KRAS*^G12D^ PDACs have higher IFNA and pSTAT3 expression, and IFNAR1 expression level inversely correlates with tumor progression. (**A**) Normalized RNA-seq reads of human IFNA13 in PDX tumor tissues; FPKM; fragments per kilobase of exon model per million reads mapped; G12D: n=15, G12V: n=10; n refers number of PDX tumor. *p<0.05 from Student’s t test. (**B-C**) Representative immunohistochemical staining (**B**) and quantification (**C**) for p-STAT3 expression in the human PDAC cells; G12D: n=5, G12V: n=5. Scalebar, left panel 1mm, right panel inserts, 250μm. *p<0.05 from Student’s t test. (**D**) IFNAR1 expression level in 5 PDCLs evaluated by Western blotting. Representative of two or more independent experiments. (**E**) Effects of exposure to recombinant (r) human IFNA (5 ng/ml) in PDCL1319 and PDCL609 *KRAS*^G12D^ cells. Total and phosphorylated (p)-STAT1 and STAT3 in cells were measured by Western blotting. Representative of at least two experimental repeats. (**F**) Validation of IFNAR1 expression by Western blotting. Representative of at least two experimental repeats. (**G-H**) Kaplan-Meier survival distributions in NSG mice bearing orthotopic tumors. 1319 sh-control: n=10; 1319 sh-IFNAR1 1: n=9; 1319 sh-IFNAR1 2: n=10 (**G**); 609 sh-control: n=10; 609 sh-IFNAR1 1: n=10; 609 sh-IFNAR1 2: n=10 (**H**). p values from Cox regression test.

Finally, we examined the correlation between OS and IFNAR1 expression in PDAC tissues by mining the TCGA database (n=178) using the GEPIA tool (*39*). Consistent with our data, tumor IFNAR1 expression levels below median associated with significantly longer OS in PDAC patients (**Fig. S11C**). These results further demonstrate that IFNAR1 expression in PDAC cells mediates a more aggressive tumor growth.

### IFNAR1 inhibition inhibits murine *Kras*^G12D^ PDAC growth in immunocompetent mice

Because downstream STAT1 and STAT3 are known to mediate immune responses (*23, 40, 41*),we next evaluated the impact of IFNAR1 inhibition in a *Kras*^G12D^ mutant murine PDAC model (AK4.4 cells) (*42, 43*). Similar to *KRAS*^G12D^ human PDCLs, AK4.4 murine PDAC cells had high IFNAR1 expression levels by Western blotting (**Fig. S12A**). Next, we generated IFNAR1-KD AK4.4 cells (**Fig. S12B**). We verified the suppressed STAT activation induced by recombinant mouse IFNA in these cells (**Fig. S12C**), similar to pharmacologic blockade with an anti-mouse IFNAR1 antibody in parental AK4.4 cells (**Fig. S12D**). Orthotopic implantation of IFNAR1-KD AK4.4 cells in immunocompetent FVB mice showed that genetic IFNAR1 inhibition significantly delayed murine PDAC growth (**Fig. S12E-F**). However, genetic IFNAR1 did not control malignant pleural effusion and, as a result, tumor growth delay did not translate into improved mouse survival as mice died primarily due to malignant pleural effusion (data not shown). As in the human PDAC models, we found that IFNAR1 inhibition did not affect tumor growth in a murine *Kras*^G12D^ PDAC model with low IFNAR1 expression level (KPC cells) (data not shown).

### Blocking IFNAR1 delays tumor growth and enhances the efficacy of immune checkpoint blockade (ICB) therapy in *Kras*^G12D^ murine PDAC

Beyond the differential PDAC cell-autonomous effects, downstream STAT3 induces immunosuppression in the tumor microenvironment, while STAT1 activation promotes anti-tumor immunity (*23, 40, 41*). In addition, IFNAR1 is expressed on malignant, stromal, and immune cells in human PDAC (**Fig. S12G**) (*45*). To reveal the effect of systemic/global blockade of IFNAR1 on tumor growth and response to immunotherapy, we examined the efficacy of anti-mouse IFNAR1 antibody treatment alone or with dual anti-PD1/CTLA4 antibody ICB therapy in the orthotopic AK4.4 murine PDAC model in immunocompetent mice. We found that anti-mouse IFNAR1 antibody alone and combined with ICB therapy, but not ICB therapy alone, significantly delayed tumor growth compared to control in this PDAC model (**Fig. 6A-B**). Moreover, combination therapy significantly increased median OS compared to control and each treatment alone (**Fig. 6C**). Of note, combination therapy significantly reduced the formation of malignant pleural effusions (**Fig. S13A**). We repeated the experiment and sacrificed the mice in a time-matched manner to examine the effects of combination treatment on target modulation and CD8^+^effector T cell infiltration. We collected tumor tissues after eight days of treatment and assessed STAT1, STAT3 and NF-κB activation (**Figs. 6D** and **S13B-D**). Western blotting analyses demonstrated suppression of both STAT3 and NF-κB in PDAC tissues after anti-IFNAR1 antibody treatment (**Fig. S13B**). In contrast, ICB alone did not affect STAT activation (**Fig. S13C**). Consistently, combination therapy suppressed the activation of STAT3 and NF-κB compared to control and ICB groups (**Figs. 6D** and **S13D**). In this model, the anti-IFNAR1 antibody treatment did not activate STAT1 via STATs cross-regulation, highlighting the more critical role of STAT3 in tumor response to anti-IFNAR1 blockade alone or combined with ICB.

**Fig. 6:**
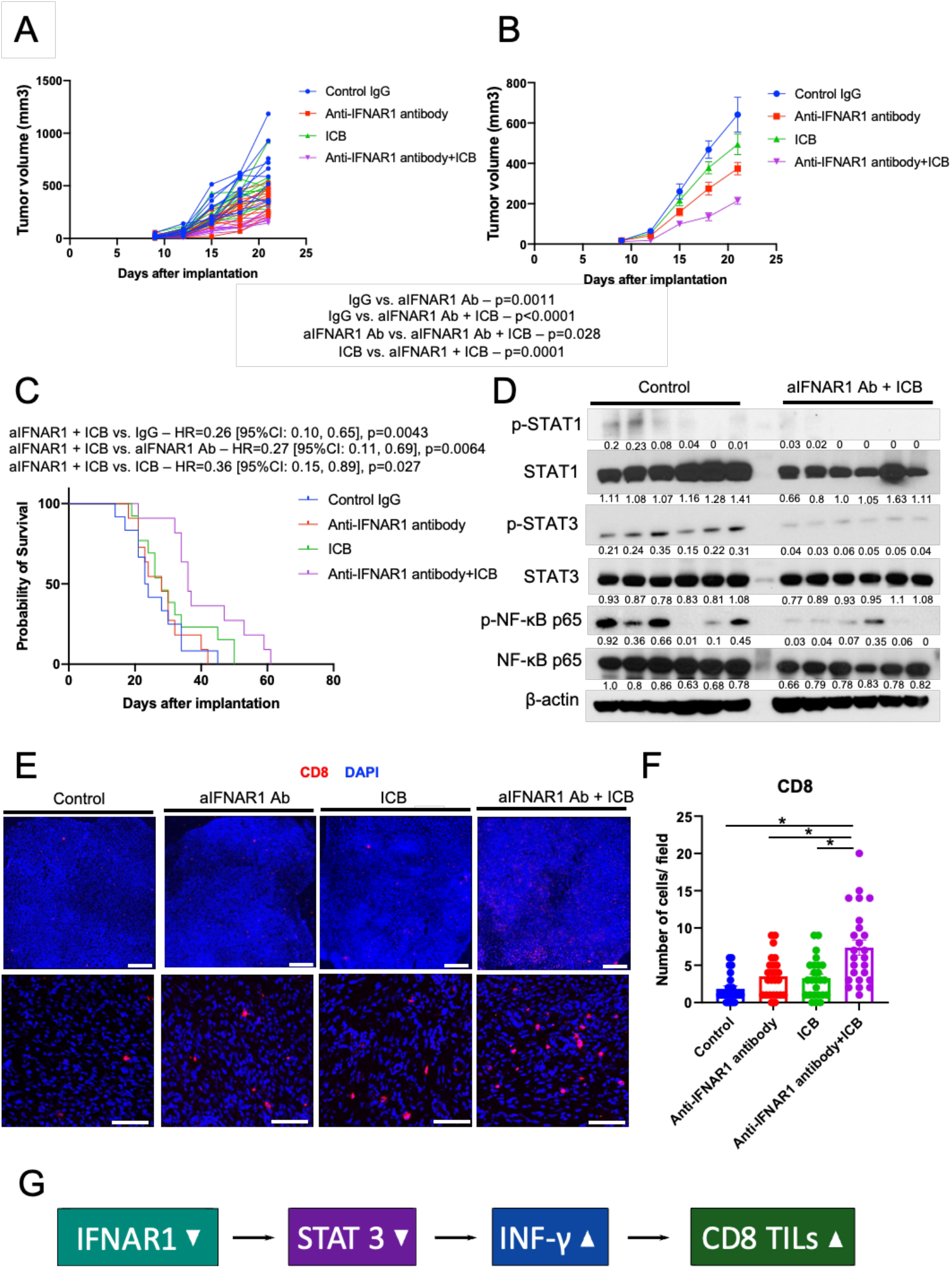
Targeting IFNAR1 renders murine PDAC responsive to immune checkpoint blockade (ICB) therapy. (**A-C**) Orthotopic tumor growth after AK4.4 implantation in FVB mice, and treatment of established tumors with either: anti-(a)IFNAR1 antibody (Ab), ICB with anti-PD1 and anti-CTLA4 antibodies, their combination, or control IgG (n=11-13 mice per group); n refers to biological replicates. Individual tumor growth curves are shown in (**A**) and average tumor size in (**B**); p value from Tukey’s test. (**C**) Kaplan-Meier survival. Distributions in the 4 treatment arms; p from Tukey’s test for tumor volume and HR from Cox regression test. (**D**) Total and phosphorylated (p)-STAT1, STAT3 and NF-κB p65 expression levels in tumor tissue. Representative of two or more independent experiments. Control: n=6; aIFNAR1 Ab + ICB: n=6. (**E**) Immunofluorescence (IF) for CD8 in tumor tissues. Scale bars are 500 μm (upper panel) and 50 μm (lower panel). (**F**) Quantification of CD8^+^ cells in tumor tissue using IF. Control IgG: n=5; a-IFNAR1 Ab: n=5; ICB: n=5; aIFNAR1Ab + ICB: n=5. *p<0.05 from Tukey’s test. (**G**) Schematic representation showing IFNAR1 blockade enhancement of immunotherapy efficacy.

Previous reports have shown that anti-IFNAR1 antibody inhibits T cell exhaustion and immunosuppression in viral infections through increased IFN-γ production (*46, 47*). Thus, we measured the IFN-γ expression level in the murine PDACs. We found increased IFN-γ levels after anti-IFNAR1 antibody treatment alone and in combination with ICB in the tumor tissues, but not after ICB alone (**Fig. S13E**). In addition, we measured the infiltration by CD8^+^ T cells in tumor tissue by immunofluorescence (IF). Consistent with the efficacy data, we found that combination therapy significantly increased the number of CD8^+^ T cells in PDAC tissue (**Fig. 6E-F**). These results show that INFAR1 blockade enhances CD8^+^ T cell infiltration and anti-PD1/CTLA4 immunotherapy efficacy in *Kras*^G12D^ murine PDAC (**Fig. 6G**).

## DISCUSSION

Although activating mutations in the *KRAS* gene are present in most PDACs, the mechanisms underlying the aggressive progression of *KRAS*^G12D^ PDACs remained obscure. In the current study, we created isogenic cell lines developed using CRISPR/Cas9 technology to reproduce the clinical behavior of these tumors and shed light on the underlying mechanisms. Surprisingly, we found that the IFNAR1/STAT3 axis, and not differential activation of canonical targets such as MAPK or PI3K pathways, mediates the more aggressive progression of *KRAS*^G12D^ versus *KRAS*^G12V^ PDACs.

IFNA belongs to the group of type I interferons, which mediate resistance to viral infections, promote antitumor activity, and modulate immune responses (*27*). Therefore, IFNA has been used as an anti-tumor drug in renal cell carcinoma and melanoma (*48*). Once secreted by cells, it binds to the same ubiquitous hetero-dimeric transmembrane receptor (IFNAR1/IFNAR2). Then, it activates canonical and non-canonical JAK/STAT signaling, which subsequently affects many genes (*49*). While most published reports showed that IFNA has anti-tumor activity, some papers revealed that IFNA could have a pro-tumor effect (*31, 50*). Since IFNA leads to cell type and context dependent patterns of interferon-stimulated gene expression via STAT modulation, IFNA might have dual functions (anti-tumor and pro-tumor) (*51*). Although some *in vivo* studies reported that rhIFNA could regulate PDAC tumor growth and enhance chemotherapy (*33, 34*), clinical trials testing combinations of IFNA with chemotherapy failed to show efficacy in unselected PDAC patient populations (*35–38*).

To determine whether and how the IFNA pathway promotes or regulates tumor progression in PDAC subsets, we conducted survival studies in loss or gain of function experiments (using genetic IFNAR1 knockdown in PDAC cells versus rhIFNA treatment). While IFNAR1 inhibition impeded PDAC progression, rhIFNA treatment showed a tendency for more aggressive tumor growth. Because STAT1 and STAT3 are downstream of IFNA signaling, we evaluated STAT1/3 activation status in these tumors. We found that STAT3 was activated while STAT1 was suppressed in *KRAS*^G12D^ versus isogenic *KRAS*^G12V^ PDACs. Since STAT1 and STAT3 may have opposing functions and have balanced expression through cross-regulation (*22, 25, 41*), we anticipated that constitutive STAT3 activation downstream IFNAR1 suppressed STAT1. Consistent with our hypothesis, when we inhibited IFNAR1, STAT3 was suppressed, and STAT1 was activated in *KRAS*^G12D^ PDAC cells. By establishing STAT3 overexpression models, we demonstrate its critical role downstream IFNAR1 as evidenced by successful tumor growth rescue studies in IFNAR1 knockdown *KRAS*^G12D^ and *KRAS*^G12V^ PDAC cells. The conclusion that STAT3 mediates IFNAR1-mediated PDAC growth is further supported by results in double knockdown for STAT1 and IFNAR1, which showed no differences in survival.

A role for the IFNA pathway and STAT3 in promoting tumor progression has been proposed in inflammatory breast cancer (IBC) (*52*). IFNA activation of STAT3 can promote anti-apoptotic processes via PI3K/AKT signaling stimulation (*53*). These studies suggested that chronic inflammation in tumors may contribute to preferential activation of STAT3 versus STAT1. PDACs are often associated with severe chronic inflammation. Indeed, we discovered that IFNAR1 blockade suppressed STAT3 and activated STAT1 in PDAC, thus delaying tumor growth, in contrast to rhIFNA treatment. Several lines of evidence support our findings. Mining of the TCGA database showed that IFNAR1 expression is negatively correlated with OS. We further demonstrate that genetic and pharmacologic inhibition of IFNAR1 delays tumor progression and is dependent on INFAR1 expression levels.

Finally, we also investigated the impact of IFNAR1 blockade on murine PDAC models in immunocompetent mice, since it has been reported that STAT1 activation enhances anti-tumor immunity and conversely, STAT3 promotes an immunosuppressive environment (*23, 40, 41*). Moreover, we tested the impact on immunotherapy since PDACs are notoriously resistant to ICB. We found that IFNAR1 blockade delayed tumor growth and enhanced the efficacy of ICB therapy. While counterintuitive, these results are supported by reports from the field of infectious diseases. Type I IFN signaling is activated by chronic virus infection and causes an immunosuppressive environment. In addition, anti-IFNAR1 antibody alleviates T cell exhaustion and immunosuppression through IFN-γ production (*46, 47*). These results indicate that anti-IFNAR1 antibody treatment can result in immune activation. Indeed, we found that IFN-γ was upregulated by anti-IFNAR1 antibody treatment and increased the number of tumor-infiltrating CD8^+^ T cells in the PDAC microenvironment.

In summary, we demonstrate that the IFNAR1/STAT3 axis is a driver of PDAC progression and mediates the more aggressive phenotype of *KRAS*^G12D^ mutant PDACs. Moreover, blockade of IFNAR1 enhanced the efficacy of immunotherapy in an aggressive *Kras*^G12D^ murine PDAC model in syngeneic mice. Since the anti-human IFNAR1 antibody anifrolumab is FDA approved for systemic lupus erythematosus and is currently under clinical development for other inflammatory disease, our results indicate that this strategy should be tested in combination with ICBs in future clinical studies in this intractable disease.

## MATERIALS AND METHODS

### Cells and cell culture

We studied low-passage PDAC patient-derived xenograft (PDX) cell lines (PDCL-1108, −1319 and −609) and a collection of 25 PDXs established in the Department of Surgery from patients treated at Massachusetts General Hospital (MGH). The murine PDAC cell line AK4.4 (*Kras*^G12D^*p53*^+/-^) was established from a tumor induced in a Ptf1-Cre/LSL-*Kras*^G12D^/*p53*^Lox/+^ mouse (*54*). The murine PDAC cell line KPC (*Kras*^G12D^ and *p53*^+/-^) was kindly provided by Dr. Saluja (Department of Surgery, University of Minnesota Medical School); it was established from a tumor induced in an LSL-Kras^G12D^/LSL-Trp53^R127H^/Pdx1-Cre mouse (*55*).

### Generation of isogenic PDCLs

To maximize genetic similarity, a cell line derived from a single-cell clone of PDCL-1108 was used for generation of CRISPR/Cas9 models. We designed spCas9 guide RNAs to target codon 12 of the human *KRAS* gene and selected the one with highest targeting efficiency determined by T7E1 mismatch assay (EnGen Mutation Detection Kit, New England Biolabs, Ipswich, MA). Truncated guide RNA was produced by PCR assembly of a guide RNA template (56, 57), followed by T7 *in vitro* transcription using the HiScribe T7 In Vitro Transcription Kit (New England Biolabs, Ipswich, MA) and purification with Trizol (ThermoFisher Scientific, Waltham, MA) according to the manufacturer’s protocols. Single-stranded oligo-deoxynucleotide (ssODN) sequences coding for the desired mutations (G12R, G12V, G12D and G12 wild type) and a silent restriction site (HindIII) for screening purposes were designed with ~ 80 bp-homology arms flanking each side of the Cas9-induced double-strand break. For transient transfection, 1 x 10^6^ PDCL-1108 cells were electroporated with 10 μg spCas9-NLS protein (New England Biolabs, Ipswich, MA), 4 μg guide RNA and 200 pmol ssODN using the Amaxa Nucleofector II (Lonza, Basel, Switzerland). Cells were recovered in growth medium for 24 hr and then sorted as single cells into 96-well plates by FACS. After colony formation, clones were screened using end-point PCR and restriction digest, followed by verification of successful editing with Sanger sequencing and targeted next-generation sequencing (CRISPR sequencing, MGH DNA core, Cambridge, MA).

### Orthotopic PDAC models in mice

We used nonobese diabetic/severe combined immunodeficiency/gamma (NSG) as well as NSG-human-HGF-knock-in (NOD.Cg- *Hgf^tm1.1HGF)Aveo^ Prkdc^scid^ Il2rg^tmlWjl/J^*) mice (Jackson Labs) for PDCLs, and FVB and C57Bl/6 mice (Jackson Labs) for AK4.4 and KPC murine PDAC cell lines, respectively. All experimental mice were bred and maintained in our gnotobiotic animal colony. All surgical procedures were performed under sterile conditions in a laminar-flow hood. Orthotopic pancreatic tumors were generated by implanting 1×10^5^ cells into the pancreas of 6-8 weeks old mice (*58*). All experimental use of animals followed the Public Health Service Policy on Humane Care of Laboratory Animals and the protocol was approved by the institutional animal care and use committee (IACUC) at MGH.

Orthotopic tumor growth and treatment responses were monitored by ultrasound imaging in mice. For survival studies, mice were monitored and euthanized when the clinical endpoint was reached, i.e., when mice became moribund. For treatment studies, we randomized mice and started treatment when tumors reached 4-5 mm in diameter. For time-matched studies, we sacrificed the mice and collected tumor tissues when the largest tumor reached 8-9 mm in diameter. Anti-mouse IFNAR1 (clone MAR1-5A3, 10 mg/kg on first dose and 5 mg/kg for the following 5 doses, every 3 days), anti-mouse CTLA-4 (clone 9D9, 10 mg/kg, 3 doses, every 3 days) and anti-mouse PD-1 antibodies (clone RMP1-14, 10 mg/kg, 6 doses, every 3 days) were purchased from BioXcell. All drug treatments were administered intraperitoneally (i.p.).

### Cell proliferation assays

To analyze viability, cells were seeded onto 96-well plates (n=8 wells). Cells were incubated at 37°C and 5% CO_2_. Cell proliferation was assessed based on the colorimetric MTT assay according to the manufacturer’s protocol. To evaluate the cell viability in 3-*D* culture conditions, we used NanoCulture Plates (ORGANOGENIX, Japan). We seeded cells (5 x 10^3^ cells/100μl) in each well under serum starvation. Cell viability was assessed using CellTiter-Glo assay (Promega, WI) 10 days after cell seeding.

### RNA sequencing analyses

Total RNA was extracted from tumor tissues using Qiagen kits. The quality control of total RNA, library preparation and sequencing were performed at the Molecular Biology Core Facilities, Dana Farber Cancer Institute (Boston, MA) with single-end 75 bp mode. Cutadapt was employed to remove the low-quality bases and adapter contamination. Next, Hisat2 was used to align the reads to the reference genome, with default alignment options and mm10 for mouse reference genome and hg38 for human reference genome (*59–61*). After mapping, samtools (*62*) was used to transfer SAM files to BAM files, and sort and build the index of BAM files. HTSeq-count (*63*) was employed to generate the count matrix. After that, edgeR (*64, 65*) was used to calculate the differentially expressed genes with cutoff of |log(fold change)| > 1 and p value < 0.001. Gene ontology functional annotation was performed by the DAVID database (https://david.ncifcrf.gov) (*66, 67*). GSEA analysis was performed using GSEA software (https://www.gsea-msigdb.org/gsea/index.jsp) with MSigDB C2 KEGG pathway and C5 Gene ontology gene sets as references.

### Quantitative real-time reverse transcription polymerase chain reaction (qPCR)

Total RNA was isolated using RNeasy Mini Kit (Qiagen Inc.) and measured by nanodrop (ThermoFisher). qPCR was performed using iTaq Universal SYBR Green Supermix (Bio-Rad, Inc.). GAPDH was used as the housekeeping gene. qPCR was done at the annealing temperature of 60°C (see **Supplemental Table S1** for primers). The relative mRNA level was calculated by the 2-ΔΔCT method.

### Protein extraction and Western blotting

The cultured cells and tissues were lysed in RIPA buffer. For immunoblotting, the cell lysates were loaded on 8% sodium dodecyl sulfate (SDS)-polyacryl-amide gels with equal amounts of protein (10 μg) per well and transferred to PVDF membranes. The membranes were blocked using 2% FBS solution in PBS for 1 hr at room temperature. Then, they were incubated with primary antibodies overnight (**Supplemental Table S1**). Signal detection was performed by Clarity Western ECL Substrate (Bio-Rad) according to the manufacturer’s instructions. These data were quantified using ImageJ (US NIH). Value indicates ratio of target protein to β-actin.

### DNA transfection and lentivirus transduction

shRNA-knockdown experiments were performed using pLKO.1 puro/neo-based lentiviruses (**Supplemental Table S2**). Briefly, 293T cells were seeded (3 x 10^5^ cells/well) in 6 well dishes 24 hr before transfection. pLKO shRNA-DNA was transfected with psPAX2 packaging and pMD2.G envelop plasmid using Fugene reagent (Promega) according to the manufacturer’s instructions. Viral supernatant was harvested 24 and 48 hr after transfection and filtered through 0.45 μm filters. PDAC cells were infected with lentivirus expressing shRNA. After 24 hr, cells were selected by puromycin/neomycin. *In vivo* and *in vitro* experiments were performed 7-10 days after infection. For the STAT3 overexpression model, we used STAT3C lentiviral plasmid and control GFP plasmid purchased from Addgene (**Supplemental Table S2**). STAT3C carries a mutation that constitutively activates STAT3. Virus was harvested and filtered as described above. Seven days after infection, GFP-positive cells were sorted by FACS.

### Immunohistochemical staining (IHC)

All sections were deparaffined with xylene and hydrated with graded alcohols. After that, for the antigen retrieval, they were boiled at 97°C in 1 mM EDTA for 20 min and cooled at RT until 37°C. Tissue sections were washed with DW and then treated serially with 3% H2O2 solution (RT, 10 min), avidin solution (RT, 15 min), and biotin solution (RT, 15 min). Sections were washed briefly with PBS after each blocking step. After washing in PBS-T (5 min x 2), sections were treated with 10% normal donkey serum (RT, 2 hr). First, the sections were stained with anti-pSTAT3 antibody (CST #9145S) at RT overnight, and then with PO-conjugated secondary antibody (Jackson #111-035-144) for 2 hr. pSTAT3 were detected using DAB-Cobalt substrate kit (Bioenno Tech #003843). Before the second staining step, sections were boiled in stripping buffer at 98°C for 20 min to inactivate the antibodies. After washing the sections with PBS and PBS-T, we treated them with 10% normal donkey serum at RT overnight. Finally, the sections were stained with anti-rodent specific COX IV antibody (CST #38563) at RT overnight and then PO-conjugated secondary antibody (Jackson #111-035-144) for 2 hr. After each antibody reaction, sections were washed with PBS-T and PBS (10 min x 3). COX IV was detected using DAB substrate kit (Abcam #ab64238). After stopping the reaction, the sections were dehydrated with graded alcohols and xylene and mounted with Malinol.

### Immunofluorescence (IF)

Tumor tissue was embedded in OCT compound, snap-frozen and cut into 6 μm thick sections.CD8^+^ T lymphocytes cells were identified by positive staining (overnight at 4°C) with anti-CD8 (Biorbyt, Saint Louis, MO) followed by incubation with Cy3-conjugated anti-rabbit antibodies (Jackson ImmunoResearch, West Grove, PA) for 2 hr at RT. Slides were prepared using ProLong^™^ Gold Antifade Mountant, and cell nuclei were identified with DAPI (Thermo Fisher Scientific, MA, Waltham, MA). All images were taken with a confocal microscope (FLUOVIEW FV1000) (OLYMPUS, Center Valley, PA). For analyses of CD8^+^ T cells, the number of cells was counted in 5 random fields under 400× magnification. These data were analyzed using ImageJ (US NIH).

### Statistical analyses

All analyses were performed using JMP Pro 11.2.0 (SAS Institute Inc., NC) and data are presented as mean ± S.E.M. Differences between experimental groups were considered statistically significant for *p*-values of less than 0.05. To compare two groups with quantitative variables, we used Student’s t test. When experimental cohort includes more than three groups with quantitative variables, we used one-way ANOVA with Tukey’s multiple comparisons test. The Kaplan-Meier method was used to generate survival curves and Cox proportional hazard model was employed to conduct comparison. Hazard ratio (HR) and 95% CI were calculated for overall survival analyses.

## List of Supplementary Materials

Materials and Methods

Fig. S1 to S13

Table S1 to S2

## Notes

## Acknowledgments

The authors thank Sylvie Roberge and Anna Khachatryan and Mark Duquette (MGH) for outstanding technical support and Drs. Cyril Benes and J. Keith Joung (MGH) for useful discussions.

## Funding

Samuel Singer Brown Fund for Pancreatic Ductal Adenocarcinoma Research (DGD)

Andrew L. Warshaw, MD Institute for Pancreatic Cancer Research Grant (DGD)

NCI Proton Beam/Federal Share Program grant (DGD)

National Institutes of Health grant R01CA247441 (DGD, LLM)

National Institutes of Health grant R01CA260872 (DGD)

National Institutes of Health grant R01CA260857 (DGD)

National Institutes of Health grant R03CA256764 (DGD)

National Institutes of Health grant U01CA224348 (RKJ)

Department of Defense grant W81XWH-19-1-0284 (DGD)

Department of Defense grant W81XWH-21-1-0738 (DGD)

Humboldt Foundation National Postdoctoral Fellowship (DHS)

Boehringer Ingelheim Fonds MD Fellowship (AK)

Fondation René TouraineFellowship (GG)

Cells in Motion cluster of excellence of Münster University Fellowship (GG)

German National Merit Foundation Fellowship (GG)

The funders had no role in the preparation of this article.

## Author contributions

Conceptualization: KI, DHS, TSH, RKJ, NB, DGD

Methodology: KI, DHS, AM, PL, SK, SA, HT, SQT, GG, AK, DF, PH, TS, CFdC, RKJ, AL, NB, DGD

Investigation: KI, DHS, AM, PL, SK, SA, HT, HK, JC, ZL, TCES, MI, GG, AK Visualization: KI, DHS, AM, PL, LLM, DGD

Funding acquisition: DHS, AK, EM, GG, LLM, DGD Project administration: DGD

Supervision: DGD

Writing – original draft: KI, HT, DGD

Writing – review & editing: All authors

## Data and materials availability

All data are available in the main text or the supplementary materials.

