## Supplemental figure, legends and tables for "IFNA pathway drives the more aggressive phenotype of *KRAS*^G12D^-mutant pancreatic ductal adenocarcinomas via IFNAR1/STAT3 activation"

#### Supplementary Figure Legends

**Fig. S1: Validation of CRISPR/Cas9 editing of *KRAS* allele.** (**A**) Determination of cleavage efficiency in PDCL-1108 cells: $1x10^{6}$ cells were transfected with sgRNA/Cas9 RNP complex and ssODN to exchange aminoacid 12R. T7E1 assay was performed 24 hr after transfection and cleavage efficiency calculated. (**B**) Knock-in efficiency in pooled 1108 cells 24 hr after transfection with Cas9/sgRNA RNP and ssODN (G12V), determined by RFLP based on introduced HindIII restriction site. Uncut wild type allele (1636 bp) and restriction fragments (855 bp and 781 bp) resulting from successful knock-in. Knock-in efficiency was calculated by dividing combined band intensities of restriction fragments with total band intensity. (**C**) Individual single- cell clones identified via elimination of a naturally occurring SacI restriction site in G12R alleles by knock-in. Non-G12R allele (1636 bp) and cut G12R restriction fragments (870 bp and 766 bp).

**Fig. S2: PDAC cells with different *KRAS* alleles have differential proliferation rates *in vitro*.** (**A**) Cell viability assays in 2-*D* using isogenic cell lines. Mean cell viability is shown as a ratio to control cells (day 1) and is plotted with error bars representing SEM (n=16). Assays were performed in duplicate; p value from Tukey's test. (**B**) Representative images of 3-*D* culture of isogenic cell lines; scale bar=320 μm. (**C**) Cell viability assays in 3-*D* using isogenic cell lines. Mean luminescence is indicated with error bars representing SEM (n=18). Assays were repeated twice; p values from Tukey's test. (**D**) Kaplan-Meier plot depicting mouse survival distributions. Isogenic human PDAC cell lines implanted in NOD-SCID-gamma (NSG) mice. G12D: n=11; G12R: n=11; G12V: n=12; Wild Type: n=12; n refers to biological replicates; p values from Cox regression analysis.

**Fig. S3: Isogenic PDAC cells with different *KRAS* alleles do not show differential activation of ERK and AKT.** (**A**) *KRAS* expression in isogenic cells with different alleles, measured by

Western blotting. Representative of two or more independent experiments. (**B**-**C**) Total and phosphorylated ERK/AKT expression *in vitro* in isogenic cells with different *KRAS* alleles, measured by Western blotting (**B**) and in tumor tissues (**C**). Representative of two or more independent experiments. (**D**) Quantification of western blot in Fig. S3C. (**D-H**) Effect of pharmacologic inhibitors of BRAF (dabrafenib and vemurafenib) (**D**), MEK (cobimetinib, trametinib and refametinib) (**E**), ERK (ravoxertinib) (**F**), and PI3K (buparlisib) (**H**) on the viability of isogenic cells with different *KRAS* alleles.

**Fig. S4. Total and phosphorylated STAT3 expression *in vitro* in PDX-derived cells with different *KRAS* alleles, measured by Western blotting.**

**Fig. S5 GO terms related to IFNA signaling are activated in *KRAS*$^{\mathbf{G12V}}$ PDAC cells.** (**A**) The top ten enriched GO biological processes of differentially expressed genes between *KRAS* G12D (n=3) and *KRAS* G12V (n=3) tumor. GO annotation was performed by Database for Annotation, Visualization and Integrated Discovery (DAVID). Graph displays category scores as $-\log_{10}$ (p value) from Hypergeometric test. Orange ones are related to STAT1 signaling. (**B**) Differentially Expressed Genes (DEGs) included in "response to virus" GO term. p value from Hypergeometric test. (**C**) Real-time qPCR validation of DEGs with tumor tissue. Mean relative mRNA level is indicated with error bars representing SEM. Assays were performed in triplicate and/or quadruplicate. n=4; n refers to biological replicates. *p<0.05, **p<0.01 from Student's t test. (**D**) Top five DEGs based on p value. Positive value of logFc means DEG is enriched in *KRAS* G12V. p value from Hypergeometric test. (**E**) TRAIL expression in tumor tissue measured by real-time qPCR. Mean relative mRNA level is indicated with error bars representing SEM. n=4. Assays were performed in triplicate and/or quadruplicate. *p<0.05 from Student's t test. (**F**) Total and phosphorylated NF-κB p65 expression level in tumor tissue. n=4. *p<0.01 from Student's t test. Representative of two or more independent experiments.

**Fig. S6 Effect of genetic IFNAR1 inhibition in isogenic PDAC cells with different KRAS alleles.** (**A**) Reduced activation of STATs by rhIFNA after IFNAR1 inhibition. Cell lysate was extracted after 15 minutes incubation with 10 ng/ml human recombinant IFNA. Total and phosphorylated STAT1 and STAT3 in cells were measured by Western blotting. Representative of two or more independent experiments. (**B, C**) Individual growth curves (**B**) and average tumor size (**C**) with SEM in surviving mice. Cancer cells were implanted into NSG-human-HGF-knock-in mice. G12D sh-control: n=8; G12D sh-IFNAR1 1: n=9; G12D sh-IFNAR1 2: n=9; G12V sh-control: n=9; G12V sh-IFNAR1 1: n=8; G12V sh-IFNAR1 2: n=7; n refers to biological replicates. p values from Tukey's test.

**Fig. S7 Effect of genetic IFNAR1 inhibition on activation of STAT1, STAT3, NF-κB, ERK and AKT in isogenic PDAC cells with different KRAS alleles.** (**A-B**) Western blotting for total and phosphorylated ERK and AKT (**A**), and quantification (**B**) after IFNAR1 knockdown. n=5; n refers to biological replicates. *p<0.05 from Student's t test. Representative of two or more independent experiments. (**C**) Expression of genes related to type I IFN signaling in sh-control and sh-IFNAR1 1 tumor tissue measured by real-time qPCR. Mean relative mRNA level is indicated with error bars representing SEM. n=5; *p<0.05 from Student's t test. Assays were performed in triplicate and/or quadruplicate. (**D-E**) Effect of pharmacologic inhibition using an IFNAR1 blocking antibody (5 μg/ml): reduced activation of STATs by, measured by Western blotting (**D**) and cell viability, measured by MTT assay (**E**). In **E**, mean cell viability is shown as a ratio to control cells (day 0), plotted with error bars representing SEM (n=8). Assays were repeated twice. *p<0.05, **p<0.01 from Student's t test. Representative of two or more independent experiments.

**Fig. S8. Effect of treatment with recombinant IFNA on *in vivo* PDAC growth.** (**A-B**) Kaplan-Meier survival distributions in NSG-human-HGF-knock-in mice bearing human PDAC with *KRAS*$^{G12D}$ (A) or *KRAS*$^{G12V}$ (B) alleles in the pancreas. Recombinant human IFNA treatment was administered at 10,000 units/mice by daily subcutaneous injections for 28 days starting when tumors were established (~100 mm$^3$ in volume); mice in the control group received vehicle injection only. G12D control: n=9; G12D IFNA: n=10; G12V control: n=7; G12V IFNA: n=8; n refers to biological replicates.

**Fig. S9. Effect of STAT3 overexpression in IFNAR1-KD PDAC models.** (**A-D**) Quantification of data shown in Fig. 4C.

**Fig. S10. Effect of STAT1 inhibition in IFNAR1-KD PDAC models.** (**A**) Validation of STAT1 and IFNAR1 expression. IFNAR1 silenced *KRAS*$^{G12D}$ cells were transfected with sh-STAT1 or sh-control. Total STAT1 and IFNAR1 in cells were measured by Western blotting. Representative of two independent experiments. (**B**) Kaplan-Meier survival distributions in NSG-human-HGF-knock-in mice bearing orthotopic human PDACs. G12D sh-IFNAR1 1/ sh-control: n=9; G12D sh-IFNAR1 1/ sh-STAT1 1: n=9; G12D sh-IFNAR1 1/ sh-STAT1 2: n=10; n refers to biological replicates. (**C**) PDAC cell viability after treatment with the STAT1 inhibitor fludarabine or the JAK1/JAK2 inhibitor ruxolitinib. Cells were treated with inhibitors for 72 hr and cell viability was assessed by MTT assay. Mean cell viability shown as a percentage of control cells. Assays were repeated twice (n=16).

**Fig. S11. Effect of genetic IFNAR1 inhibition in PDCLs with different KRAS alleles.** (**A**) Human IFNA1 and –13 expression in PDX-derived cell lines assessed by real-time qPCR. Mean relative mRNA level is indicated with error bars representing SEM. Assay was performed in quadruplicate. *p<0.05, **p<0.01 from Tukey's test. (**B**) Reduced STATs activation by IFNAR1 inhibition in PDX-derived cell lines. Total and phosphorylated STAT1 and STAT3 in cells were measured by Western blotting. Cells were treated with 10 ng/ml recombinant human IFNA. Representative of two or more independent experiments. (**C**) Gene Expression Profiling Interactive Analysis (GEPIA) data showing correlation between IFNAR1 expression and survival in pancreatic cancer patients. Cut-off value: median, p value from Log-rank test.

**Fig. S12. Effect of genetic IFNAR1 inhibition on murine PDAC growth in immunocompetent mice.** (**A**) IFNAR1 expression in AK4.4 and KPC murine *Kras*$^{G12D}$ PDAC cells. Representative of two or more independent experiments. (**B**) Validation of genetic IFNAR1 knockdown. Representative of two or more independent experiments. (**C-D**) Reduced activation of STATs by IFNAR1 inhibition. Cell lysate was extracted after 15 minutes incubation with 10 ng/ml mouse recombinant IFNA. Cells were treated with 5 ug/ml anti-mouse (a)IFNAR1 antibody (Ab) for 24 hr. Total and phosphorylated STAT1 and STAT3 in cells were measured by western blotting. Representative of two or more independent experiments. (**E**) Individual growth curves and average tumor size with SEM in surviving mice. AK4.4 murine PDAC cells were implanted into FVB mice. AK4.4 sh-control: n=11; AK4.4 sh-IFNAR1 1: n=11; AK4.4 sh-IFNAR1 2: n=12; n refers to biological replicates. p values from Tukey’s test. (**F**) Kaplan-Meier curve depicting mouse survival. (**G**) IFNAR1 expression in human PDAC cells from Tumor Immune Single-cell Hub (TISCH) search engine using published data sets.

**Fig. S13. Effect of anti (a)IFNAR1 antibody (Ab), immune checkpoint blockade (ICB), their combination or control treatment on pleural ascites formation and on STAT1, STAT3 and NF-κB p65 activation and IFN-γ expression in murine *Kras*$^{G12D}$ PDAC.** (**A**) Number of mice with malignant pleural effusion at the experimental endpoint in the survival study (see **Fig. 6C**). (**B-D**) Total and phosphorylated STAT1, STAT3 and NF-κB p65 expression in tumor tissues after aIFNAR1 Ab (**B**), ICB (**C**) or their combination (**D**). **B**-**D**, n=6; n refers to biological replicates. Representative of two or more independent experiments. (**E**) IFN-γ expression in tumor tissue measured by real-time qPCR after anti (a)IFNAR1 antibody treatment, immune checkpoint blockade (ICB), their combination or control treatment. Mean relative mRNA level is indicated with error bars representing SEM. n=5. Assays were performed in triplicate and/or quadruplicate. **p<0.01, *p<0.05 from Student's t test.

Fig. S1

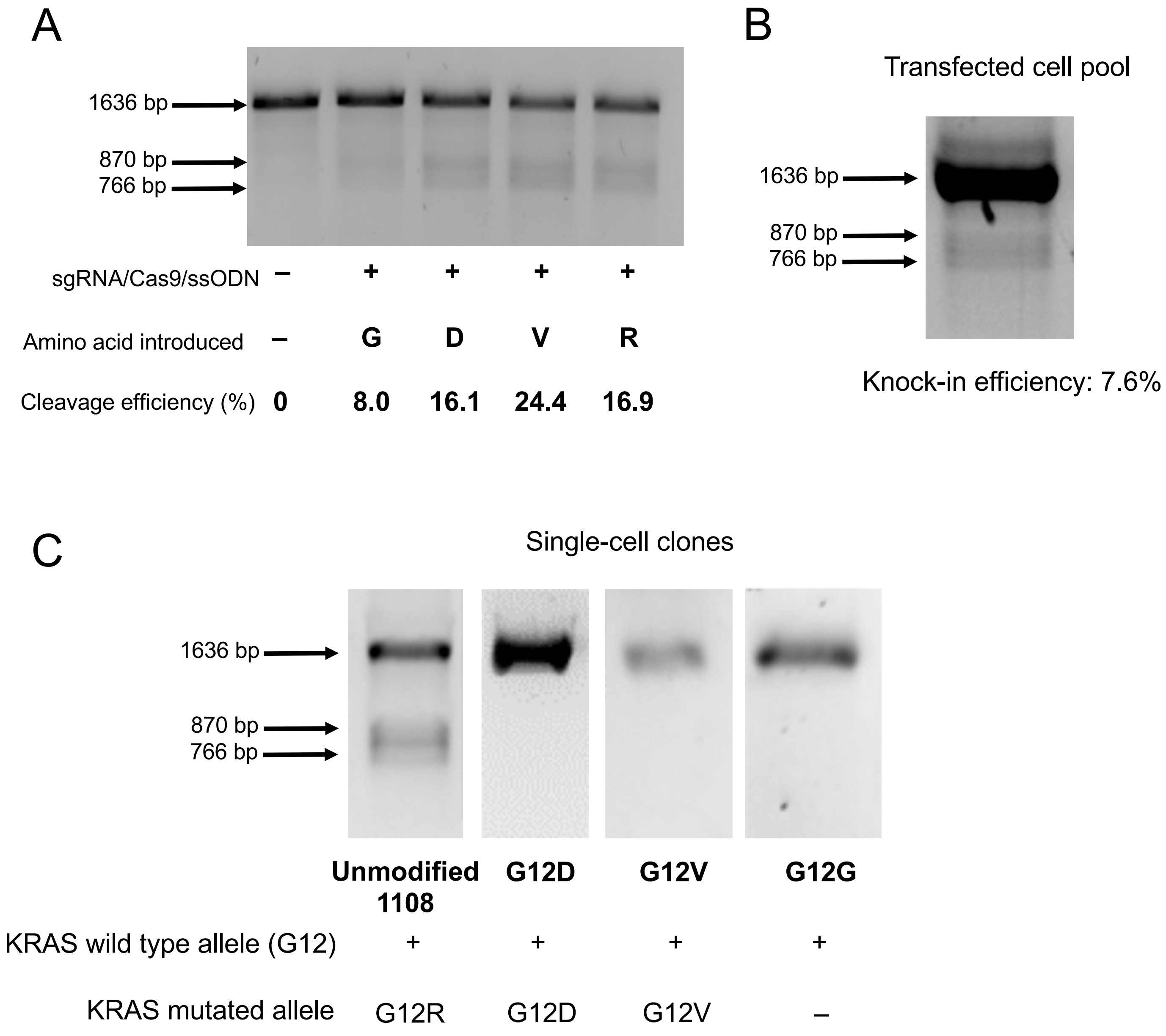

A
1636 bp
870 bp
766 bp
sgRNA/Cas9/ssODN – + + + +
Amino acid introduced – G D V R
Cleavage efficiency (%) 0 8.0 16.1 24.4 16.9
B
Transfected cell pool
1636 bp
870 bp
766 bp
Knock-in efficiency: 7.6%
C
Single-cell clones
1636 bp
870 bp
766 bp
Unmodified 1108
G12D
G12V
G12G
KRAS wild type allele (G12) + + + +
KRAS mutated allele G12R G12D G12V –

Fig. S2

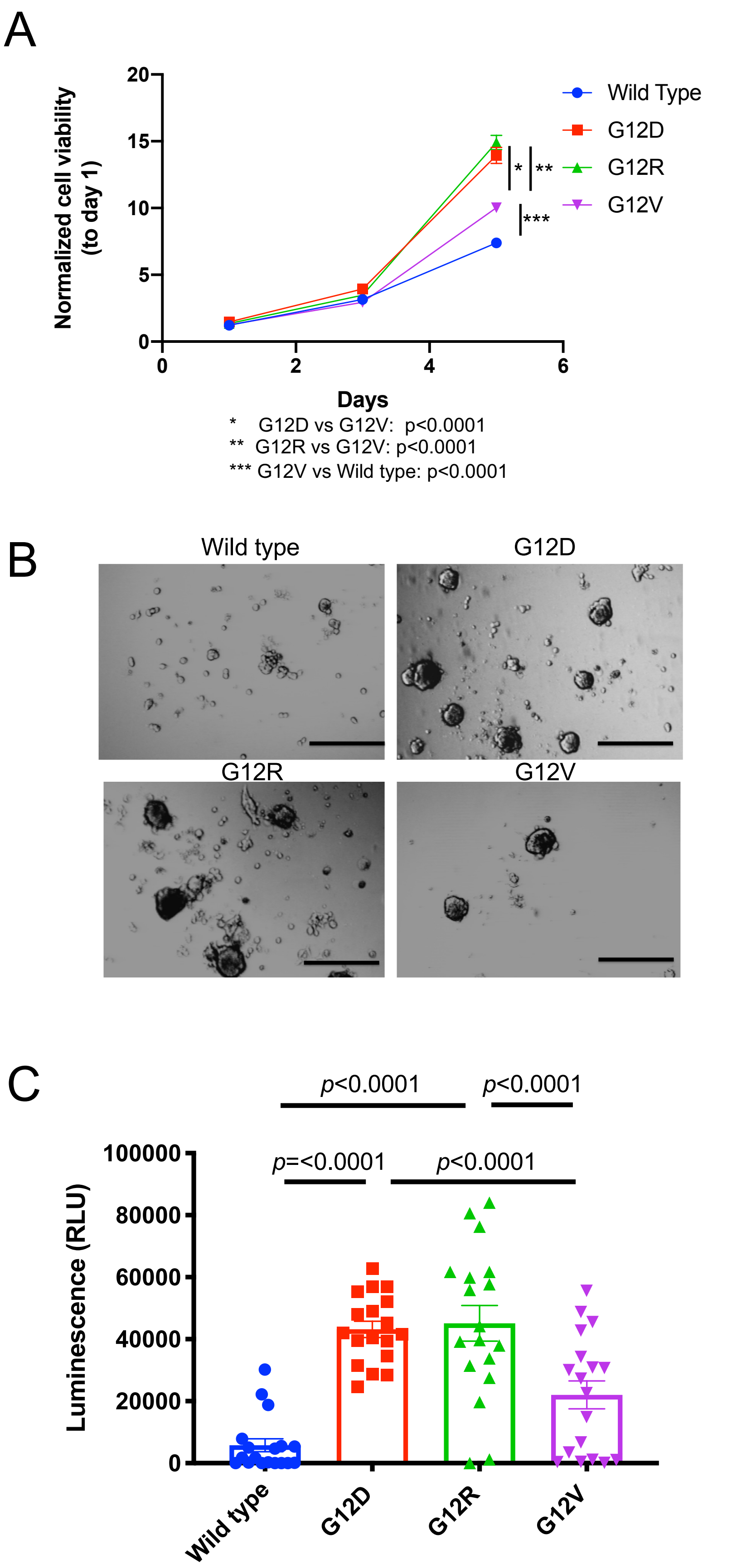
A
Normalized cell viability
(to day 1)
20
15
10
5
0
0
2
4
6
Days
Wild Type
G12D
G12R
G12V
*
**
***
*    G12D vs G12V:  p<0.0001
**  G12R vs G12V: p<0.0001
*** G12V vs Wild type: p<0.0001
B
Wild type
G12D
G12R
G12V
C
p<0.0001
p<0.0001
p=<0.0001
p<0.0001
Luminescence (RLU)
100000
80000
60000
40000
20000
0
Wild type
G12D
G12R
G12V

### Fig. S3

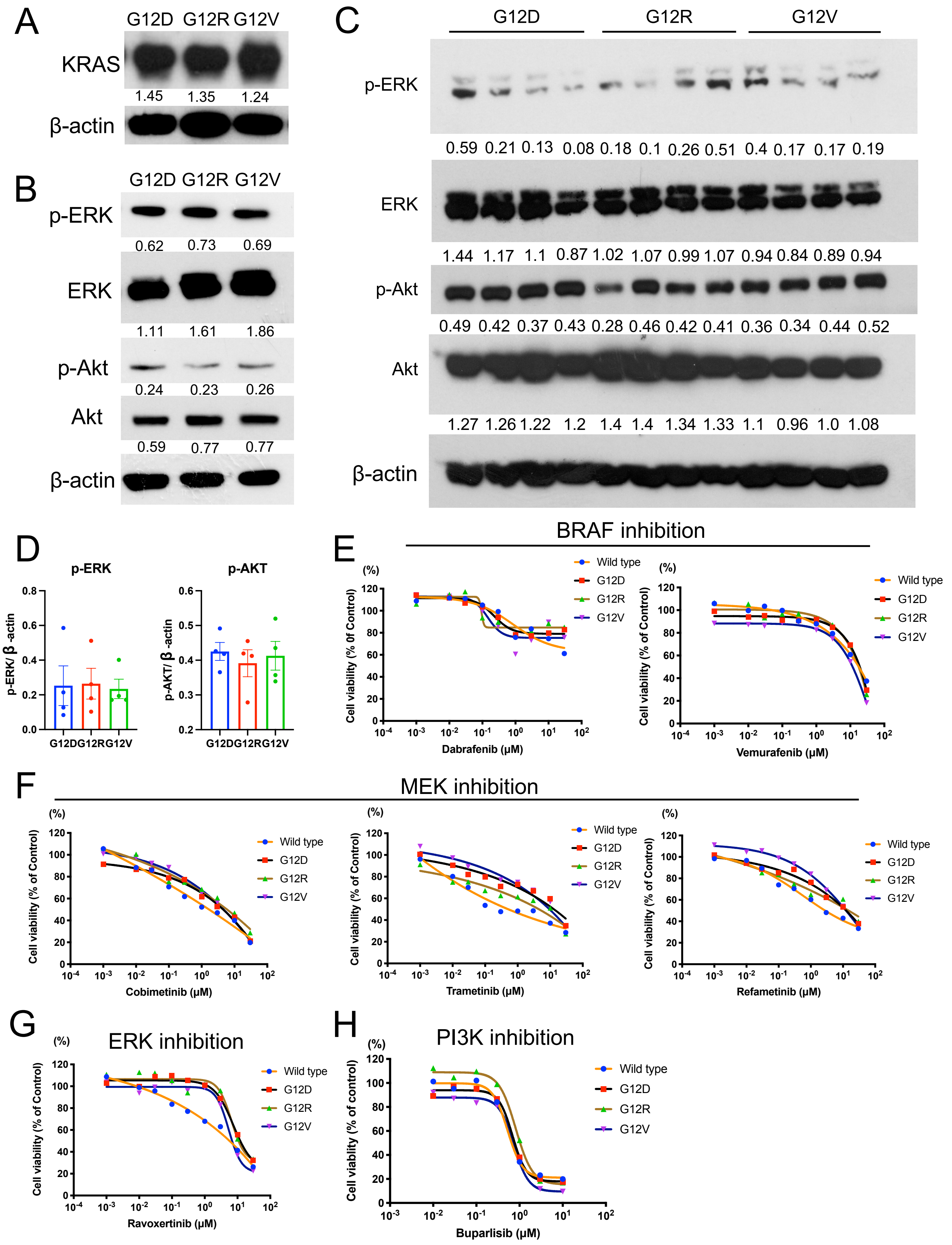
A
G12D G12R G12V
KRAS
1.45 1.35 1.24
β-actin
B
G12D G12R G12V
p-ERK
0.62 0.73 0.69
ERK
1.11 1.61 1.86
p-Akt
0.24 0.23 0.26
Akt
0.59 0.77 0.77
β-actin
C
G12D G12R G12V
p-ERK
0.59 0.21 0.13 0.08 0.18 0.1 0.26 0.51 0.4 0.17 0.17 0.19
ERK
1.44 1.17 1.1 0.87 1.02 1.07 0.99 1.07 0.94 0.84 0.89 0.94
p-Akt
0.49 0.42 0.37 0.43 0.28 0.46 0.42 0.41 0.36 0.34 0.44 0.52
Akt
1.27 1.26 1.22 1.2 1.4 1.4 1.34 1.33 1.1 0.96 1.0 1.08
β-actin
D
p-ERK
p-ERK/ β-actin
p-AKT
p-AKT/ β-actin
G12D G12R G12V
E
BRAF inhibition
Cell viability (% 0f Control)
(%)
Dabrafenib (μM)
Vemurafenib (μM)
Wild type
G12D
G12R
G12V
F
MEK inhibition
Cell viability (% of Control)
Cobimetinib (μM)
Trametinib (μM)
Refametinib (μM)
G
ERK inhibition
Ravoxertinib (μM)
H
PI3K inhibition
Cell viability (% of control)
Buparlisib (μM)

### Fig. S4

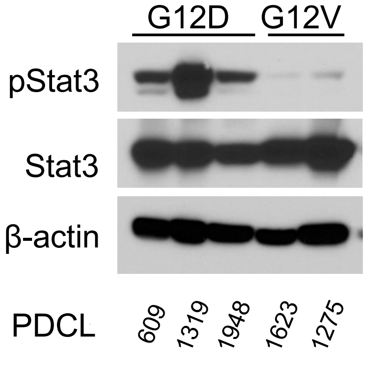

### Fig. S5

A

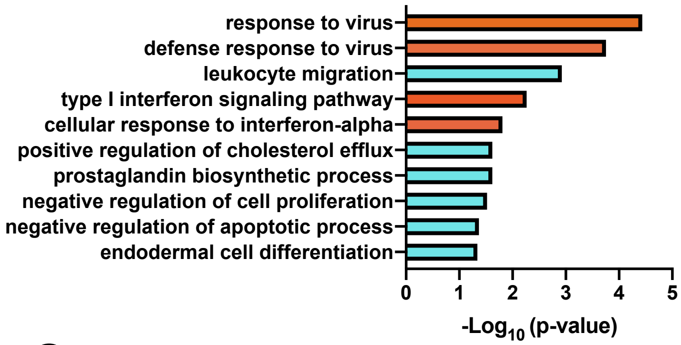


B Differentially Expressed Genes in gene set “response to virus”

| Gene | logFC | p value | FDR |
|---|---|---|---|
| IFIT3 | 1.17 | 8.58E-16 | 1.39E-12 |
| IFIT2 | 1.13 | 4.65E-11 | 1.93E-08 |
| OASL | 1.29 | 5.19E-11 | 2.01E-08 |
| TRIM22 | 1.14 | 3.09E-10 | 8.67E-08 |
| DDX60 | 1.30 | 6.71E-09 | 1.11E-06 |

logFc; positive value: enriched in *KRAS* G12V

C

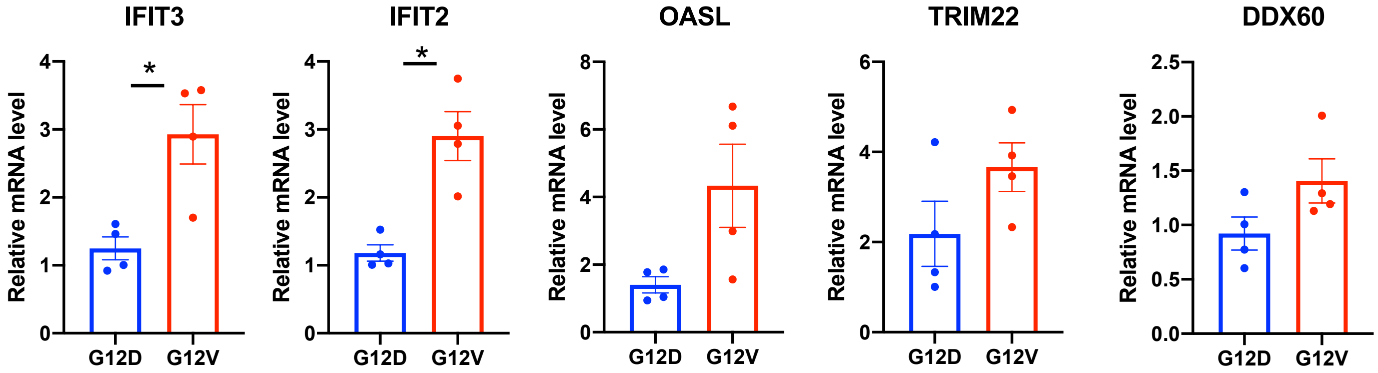


D

Top five Differentially Expressed Genes

| Gene | logFc | p value | FDR |
|---|---|---|---|
| DEGs (mapping to human) | | | |
| DUSP1 | 1.25 | 1.05E-25 | 8.49E-22 |
| TRAIL | 1.17 | 1.17E-16 | 4.74E-13 |
| GNG4 | -1.13 | 3.97E-16 | 8.73E-13 |
| GBP4 | 1.61 | 4.30E-16 | 8.73E-13 |
| IFIT3 | 1.17 | 8.58E-16 | 1.39E-12 |

logFc; positive value: enriched in *KRAS* G12V

E

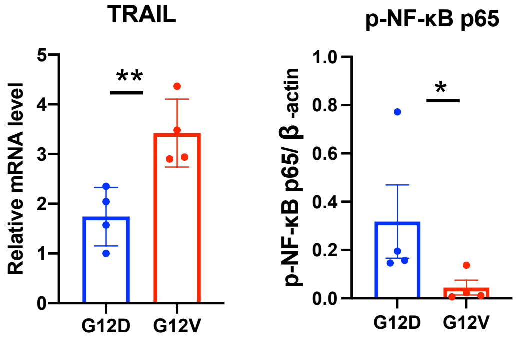


F

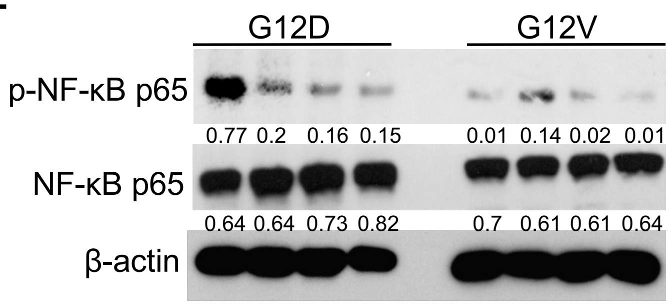

### Fig. S6

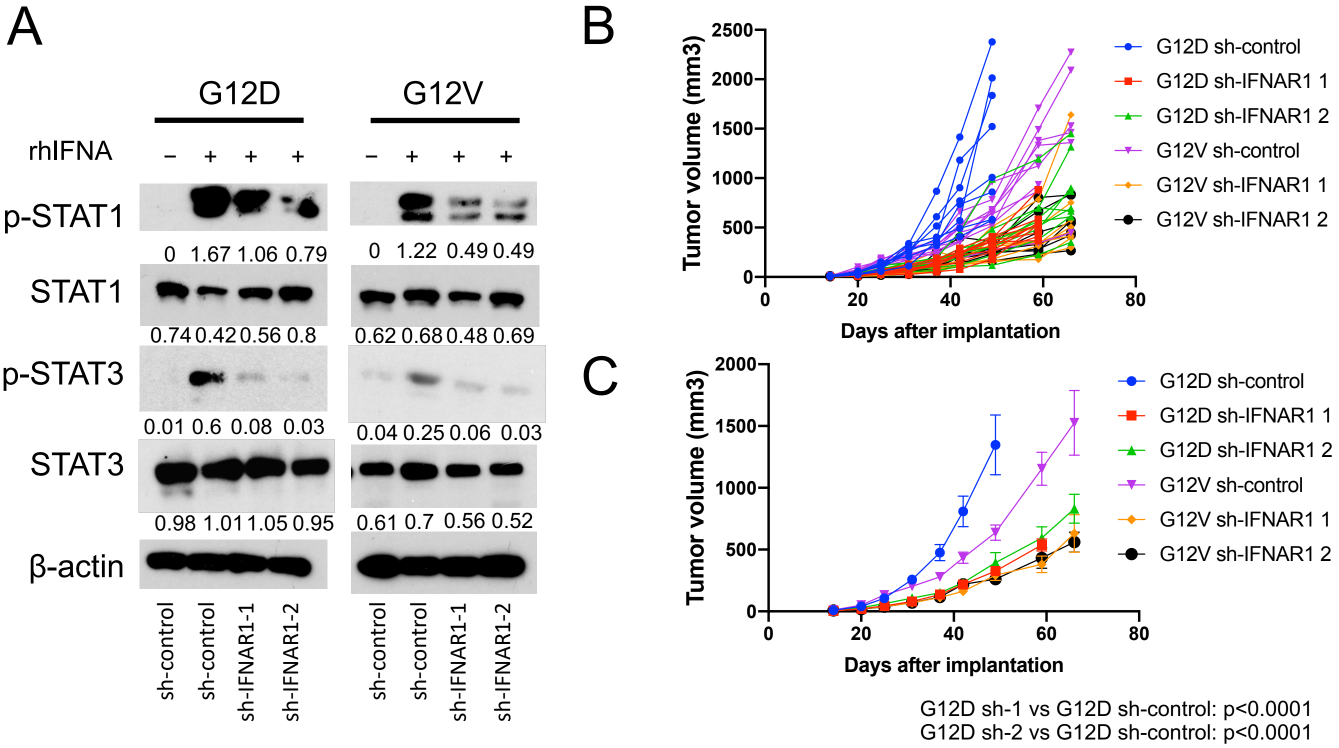


G12D sh-1 vs G12D sh-control: $p<0.0001$
G12D sh-2 vs G12D sh-control: $p<0.0001$
G12V sh-1 vs G12V sh-control: $p<0.0001$
G12V sh-2 vs G12V sh-control: $p<0.0001$
G12V sh-control vs G12D sh-control: $p=0.0004$

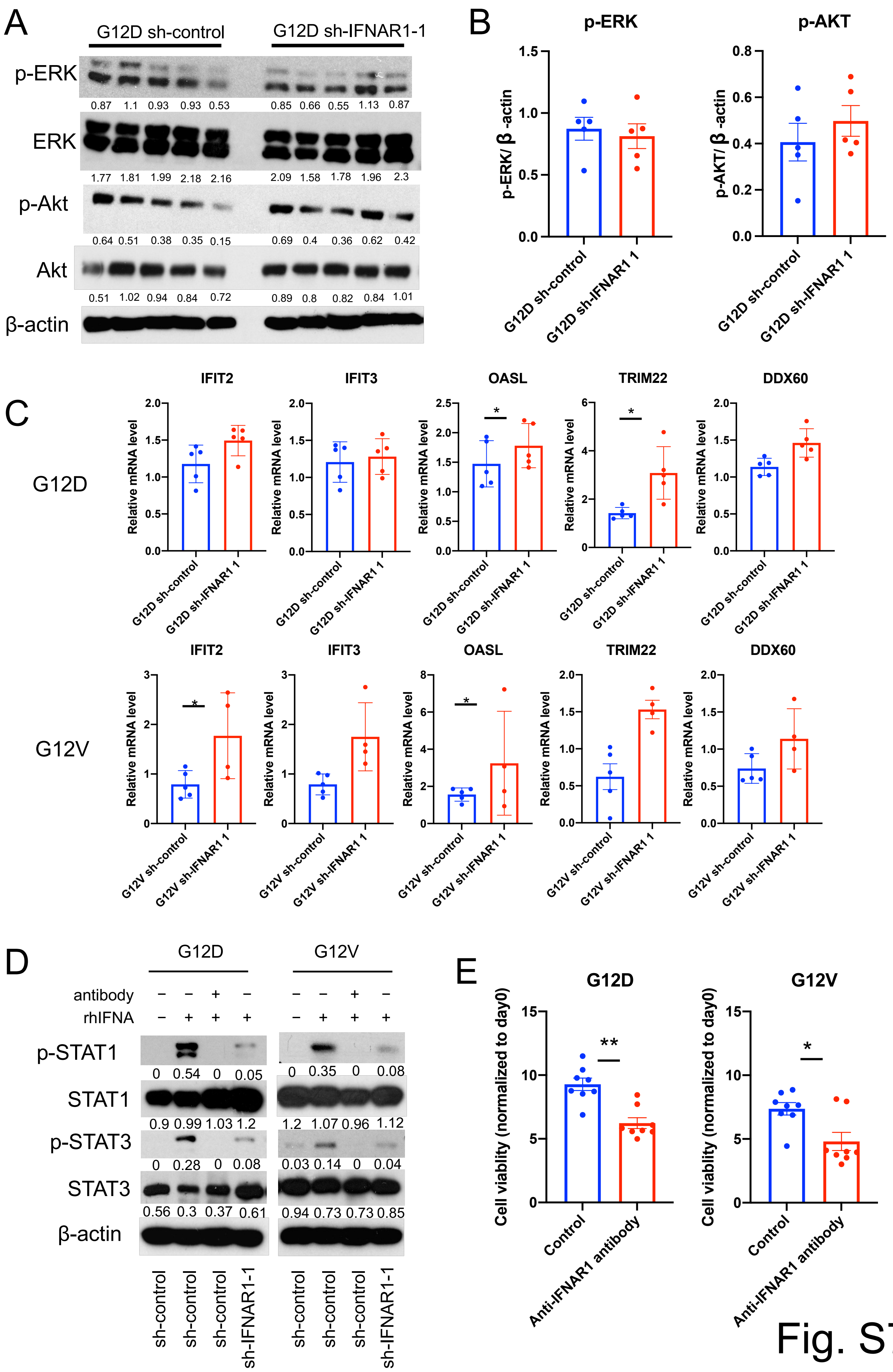
A
G12D sh-control
G12D sh-IFNAR1-1
p-ERK
0.87 1.1 0.93 0.93 0.53
0.85 0.66 0.55 1.13 0.87
ERK
1.77 1.81 1.99 2.18 2.16
2.09 1.58 1.78 1.96 2.3
p-Akt
0.64 0.51 0.38 0.35 0.15
0.69 0.4 0.36 0.62 0.42
Akt
0.51 1.02 0.94 0.84 0.72
0.89 0.8 0.82 0.84 1.01
β-actin
B
p-ERK
p-AKT
p-ERK/ β -actin
p-AKT/ β -actin
G12D sh-control
G12D sh-IFNAR1 1
C
G12D
G12V
IFIT2
IFIT3
OASL
TRIM22
DDX60
Relative mRNA level
G12D sh-control
G12D sh-IFNAR1 1
G12V sh-control
G12V sh-IFNAR1 1
D
G12D
G12V
antibody
rhIFNA
p-STAT1
0 0.54 0 0.05
0 0.35 0 0.08
STAT1
0.9 0.99 1.03 1.2
1.2 1.07 0.96 1.12
p-STAT3
0 0.28 0 0.08
0.03 0.14 0 0.04
STAT3
0.56 0.3 0.37 0.61
0.94 0.73 0.73 0.85
β-actin
sh-control
sh-control
sh-control
sh-IFNAR1-1
E
G12D
G12V
Cell viablity (normalized to day0)
Control
Anti-IFNAR1 antibody

Fig. S7

### Fig. S8

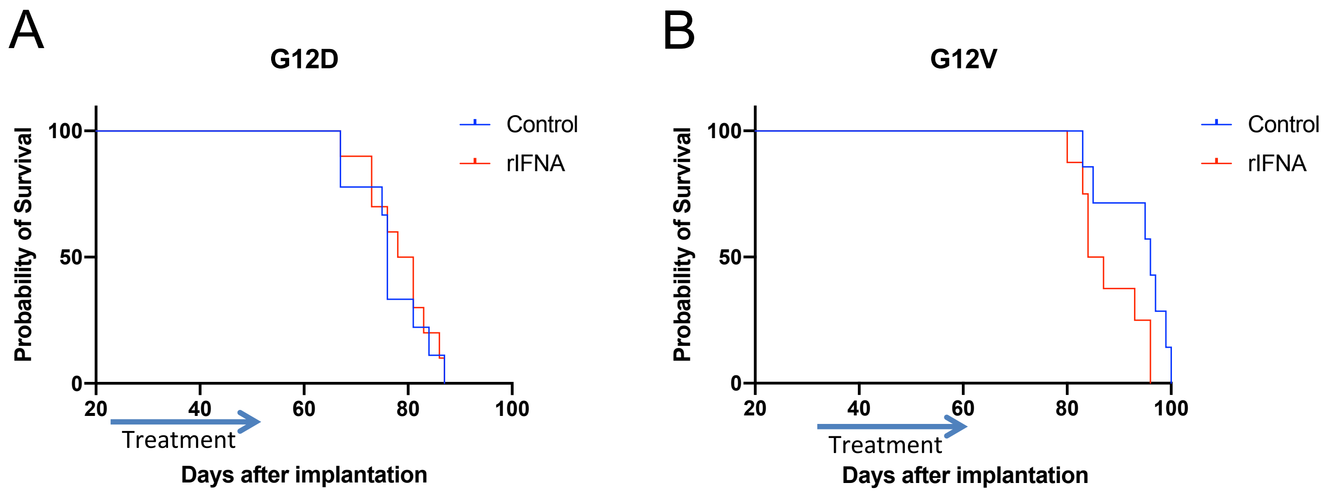


Control vs rIFNA – HR=0.443 [95%CI: 0.152, 1.293], P=0.061

Fig. S9

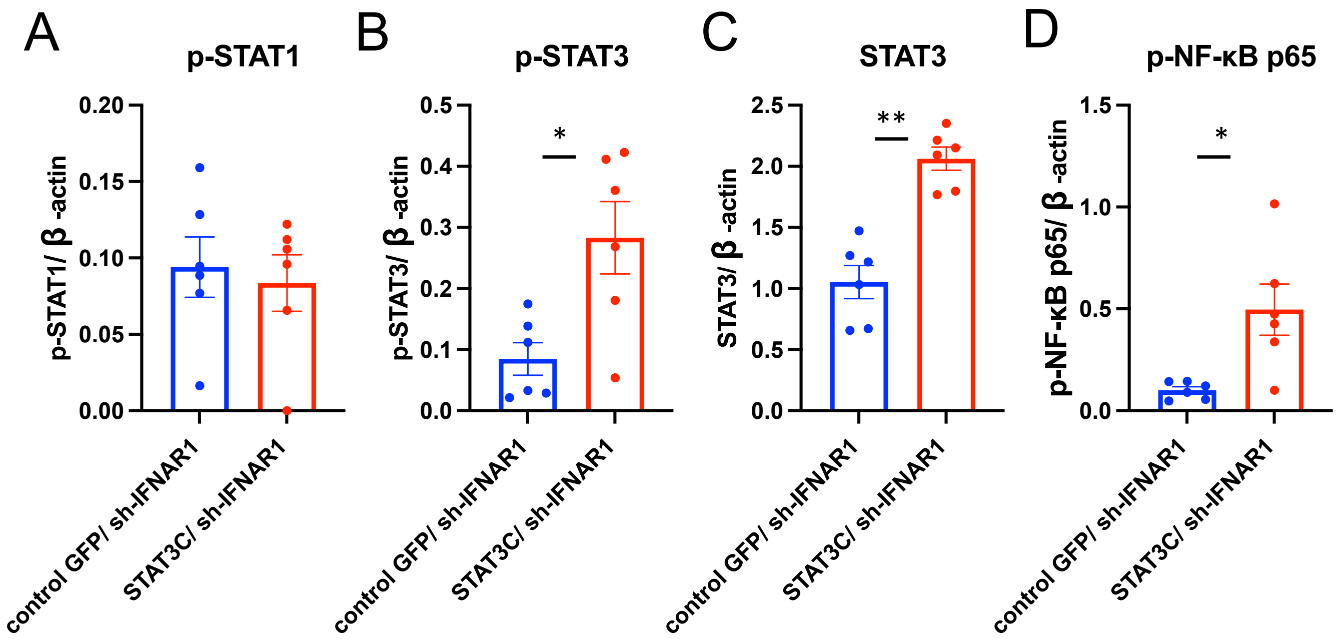
A
p-STAT1
p-STAT1/ β -actin
0.20
0.15
0.10
0.05
0.00
control GFP/ sh-IFNAR1
STAT3C/ sh-IFNAR1
B
p-STAT3
p-STAT3/ β -actin
0.5
0.4
0.3
0.2
0.1
0.0
*
control GFP/ sh-IFNAR1
STAT3C/ sh-IFNAR1
C
STAT3
STAT3/ β -actin
2.5
2.0
1.5
1.0
0.5
0.0
**
control GFP/ sh-IFNAR1
STAT3C/ sh-IFNAR1
D
p-NF-κB p65
p-NF-κB p65/ β -actin
1.5
1.0
0.5
0.0
*
control GFP/ sh-IFNAR1
STAT3C/ sh-IFNAR1

### Fig. S10

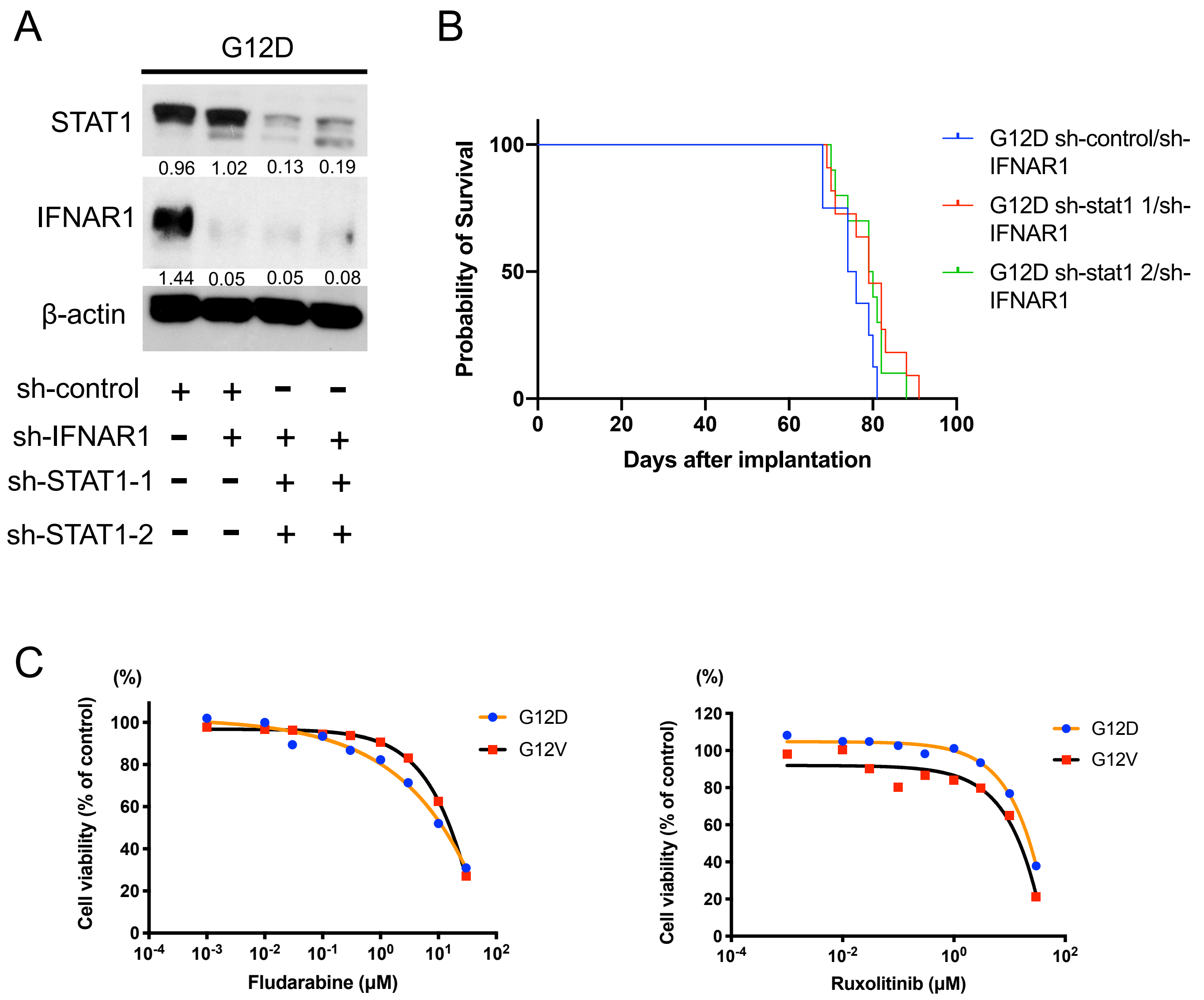
A
G12D
STAT1
0.96 1.02 0.13 0.19
IFNAR1
1.44 0.05 0.05 0.08
β-actin
sh-control + + − −
sh-IFNAR1 − + + +
sh-STAT1-1 − − + +
sh-STAT1-2 − − + +
B
Probability of Survival
100
50
0
0 20 40 60 80 100
Days after implantation
G12D sh-control/sh-IFNAR1
G12D sh-stat1 1/sh-IFNAR1
G12D sh-stat1 2/sh-IFNAR1
C
(%)
Cell viability (% of control)
0 20 40 60 80 100
Fludarabine (μM)
G12D
G12V
(%)
Cell viability (% of control)
0 20 40 60 80 100 120
Ruxolitinib (μM)
G12D
G12V

### Fig. S11

A

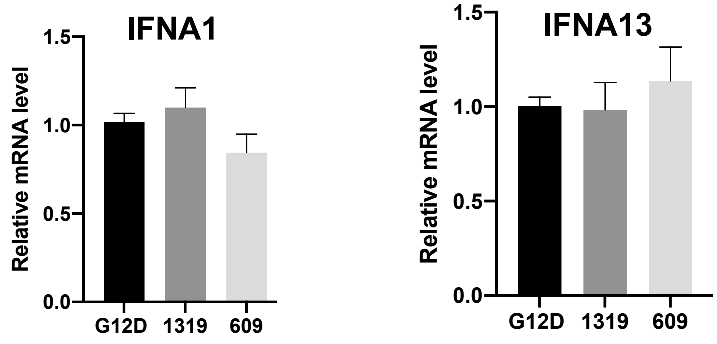
IFNA1
IFNA13
Relative mRNA level
1.5
1.0
0.5
0.0
G12D
1319
609

B

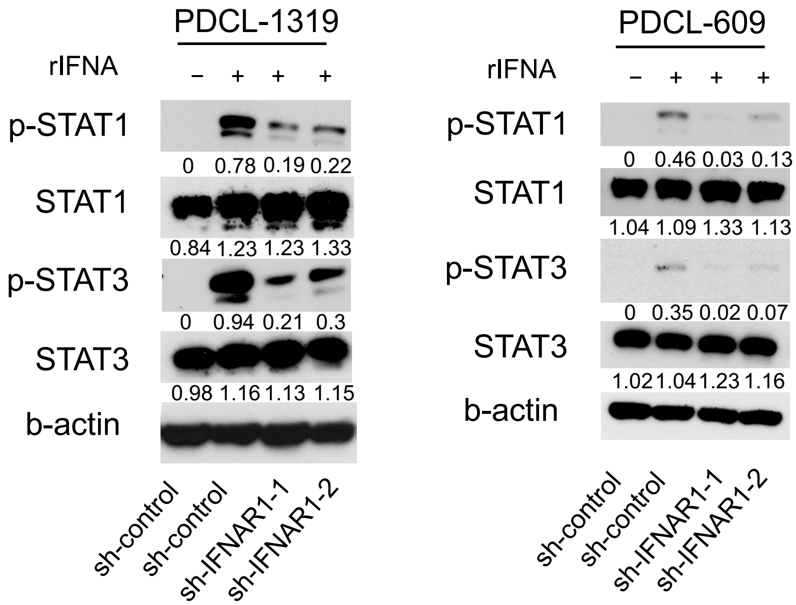
PDCL-1319
PDCL-609
rIFNA
– + + +
p-STAT1
0 0.78 0.19 0.22
0 0.46 0.03 0.13
STAT1
0.84 1.23 1.23 1.33
1.04 1.09 1.33 1.13
p-STAT3
0 0.94 0.21 0.3
0 0.35 0.02 0.07
STAT3
0.98 1.16 1.13 1.15
1.02 1.04 1.23 1.16
b-actin
sh-control
sh-control
sh-IFNAR1-1
sh-IFNAR1-2

C

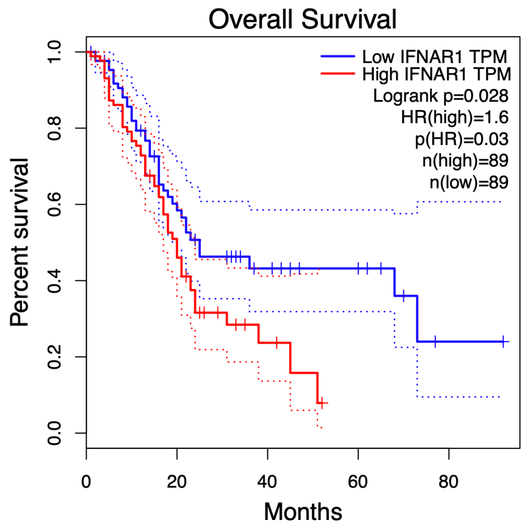
Overall Survival
Low IFNAR1 TPM
High IFNAR1 TPM
Logrank p=0.028
HR(high)=1.6
p(HR)=0.03
n(high)=89
n(low)=89
Percent survival
0.0
0.2
0.4
0.6
0.8
1.0
0
20
40
60
80
Months

Fig. S12

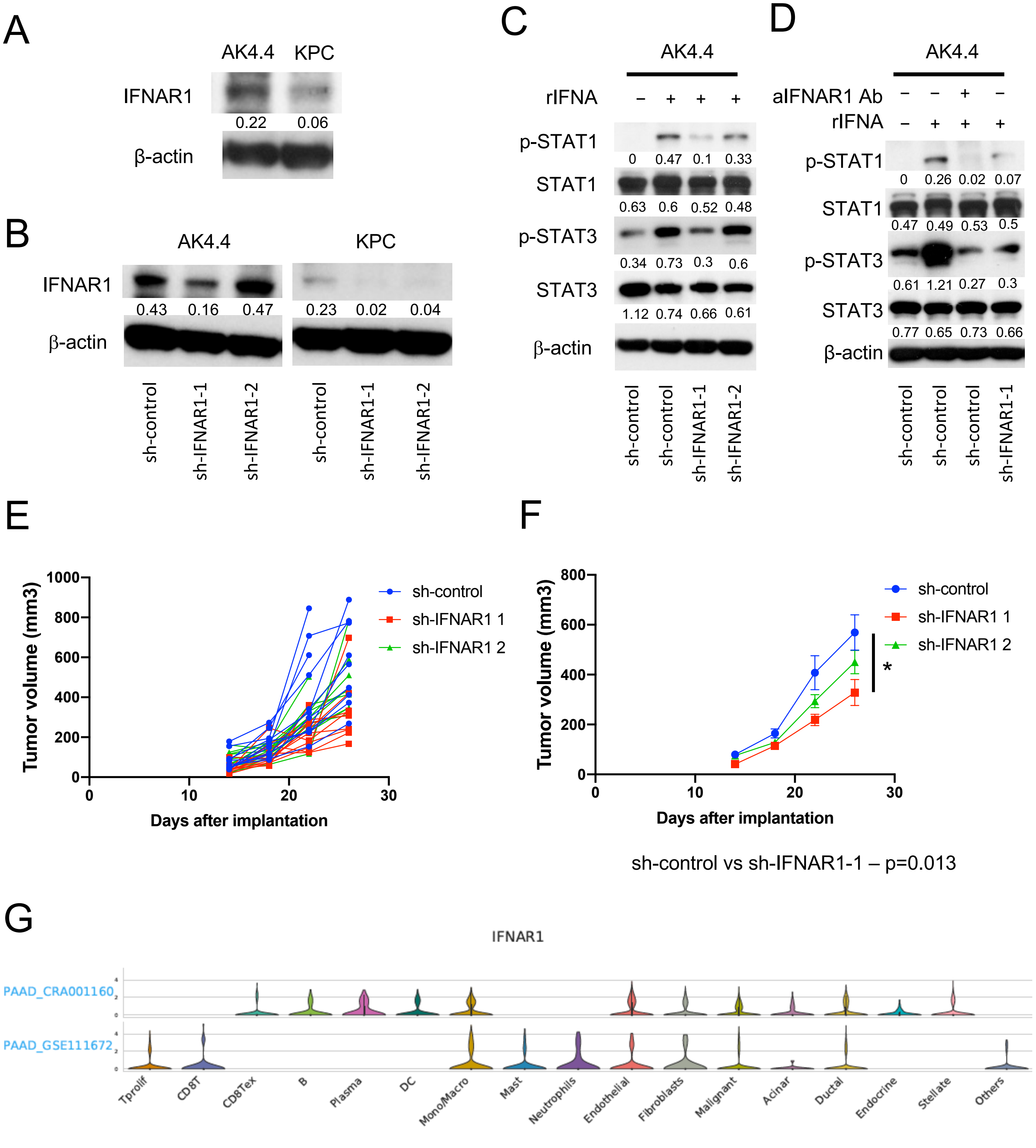
A
AK4.4
KPC
IFNAR1
0.22
0.06
β-actin
B
AK4.4
KPC
IFNAR1
0.43
0.16
0.47
0.23
0.02
0.04
β-actin
sh-control
sh-IFNAR1-1
sh-IFNAR1-2
sh-control
sh-IFNAR1-1
sh-IFNAR1-2
C
AK4.4
rIFNA
– + + +
p-STAT1
0 0.47 0.1 0.33
STAT1
0.63 0.6 0.52 0.48
p-STAT3
0.34 0.73 0.3 0.6
STAT3
1.12 0.74 0.66 0.61
β-actin
sh-control
sh-control
sh-IFNAR1-1
sh-IFNAR1-2
D
AK4.4
aIFNAR1 Ab
– – + –
rIFNA
– + + +
p-STAT1
0 0.26 0.02 0.07
STAT1
0.47 0.49 0.53 0.5
p-STAT3
0.61 1.21 0.27 0.3
STAT3
0.77 0.65 0.73 0.66
β-actin
sh-control
sh-control
sh-control
sh-IFNAR1-1
E
Tumor volume (mm3)
Days after implantation
sh-control
sh-IFNAR1 1
sh-IFNAR1 2
F
Tumor volume (mm3)
Days after implantation
sh-control
sh-IFNAR1 1
sh-IFNAR1 2
*
sh-control vs sh-IFNAR1-1 – p=0.013
G
IFNAR1
PAAD_CRA001160
PAAD_GSE111672
Tprolif
CD8T
CD8Tex
B
Plasma
DC
Mono/Macro
Mast
Neutrophils
Endothelial
Fibroblasts
Malignant
Acinar
Ductal
Endocrine
Stellate
Others

Fig. S13

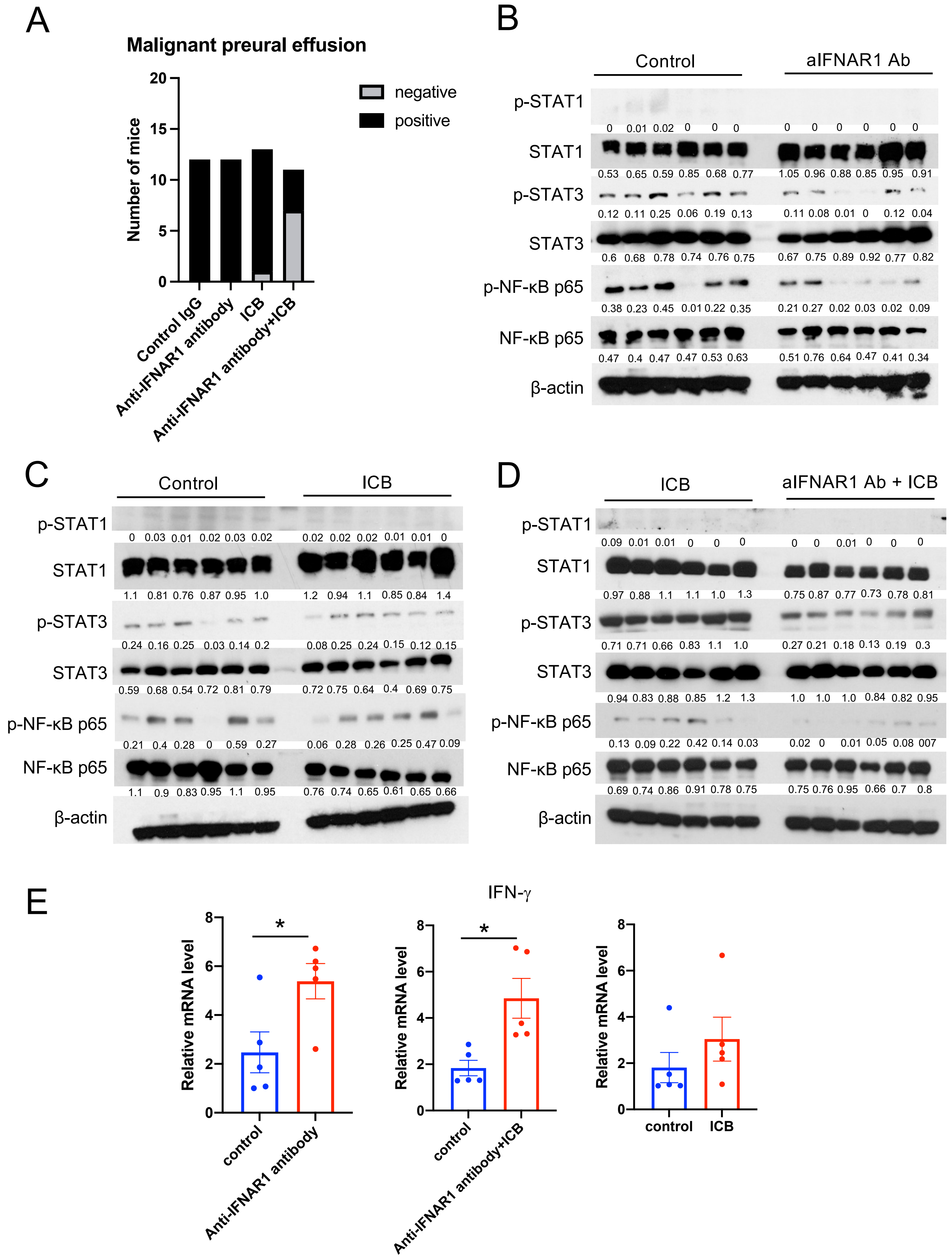
A
Malignant preural effusion
negative
positive
Number of mice
0
5
10
15
20
Control IgG
Anti-IFNAR1 antibody
ICB
Anti-IFNAR1 antibody+ICB
B
Control
aIFNAR1 Ab
p-STAT1
0 0.01 0.02 0 0 0
0 0 0 0 0 0
STAT1
0.53 0.65 0.59 0.85 0.68 0.77
1.05 0.96 0.88 0.85 0.95 0.91
p-STAT3
0.12 0.11 0.25 0.06 0.19 0.13
0.11 0.08 0.01 0 0.12 0.04
STAT3
0.6 0.68 0.78 0.74 0.76 0.75
0.67 0.75 0.89 0.92 0.77 0.82
p-NF-κB p65
0.38 0.23 0.45 0.01 0.22 0.35
0.21 0.27 0.02 0.03 0.02 0.09
NF-κB p65
0.47 0.4 0.47 0.47 0.53 0.63
0.51 0.76 0.64 0.47 0.41 0.34
β-actin
C
Control
ICB
p-STAT1
0 0.03 0.01 0.02 0.03 0.02
0.02 0.02 0.02 0.01 0.01 0
STAT1
1.1 0.81 0.76 0.87 0.95 1.0
1.2 0.94 1.1 0.85 0.84 1.4
p-STAT3
0.24 0.16 0.25 0.03 0.14 0.2
0.08 0.25 0.24 0.15 0.12 0.15
STAT3
0.59 0.68 0.54 0.72 0.81 0.79
0.72 0.75 0.64 0.4 0.69 0.75
p-NF-κB p65
0.21 0.4 0.28 0 0.59 0.27
0.06 0.28 0.26 0.25 0.47 0.09
NF-κB p65
1.1 0.9 0.83 0.95 1.1 0.95
0.76 0.74 0.65 0.61 0.65 0.66
β-actin
D
ICB
aIFNAR1 Ab + ICB
p-STAT1
0.09 0.01 0.01 0 0 0
0 0 0.01 0 0 0
STAT1
0.97 0.88 1.1 1.1 1.0 1.3
0.75 0.87 0.77 0.73 0.78 0.81
p-STAT3
0.71 0.71 0.66 0.83 1.1 1.0
0.27 0.21 0.18 0.13 0.19 0.3
STAT3
0.94 0.83 0.88 0.85 1.2 1.3
1.0 1.0 1.0 0.84 0.82 0.95
p-NF-κB p65
0.13 0.09 0.22 0.42 0.14 0.03
0.02 0 0.01 0.05 0.08 007
NF-κB p65
0.69 0.74 0.86 0.91 0.78 0.75
0.75 0.76 0.95 0.66 0.7 0.8
β-actin
E
IFN-γ
Relative mRNA level
0
2
4
6
8
*
control
Anti-IFNAR1 antibody
Relative mRNA level
*
control
Anti-IFNAR1 antibody+ICB
Relative mRNA level
control
ICB

### Supplemental Table S1: Antibody and PCR primer information

| Antibody | Company | Catalog # | Host | Dilution |
|---|---|---|---|---|
| Immunofluorescence (IF) | | | | |
| CD8 | Biorbyt | orb348907 | Rabbit | 1;100 |
| CD11b | BioLegend | 101202 | Rat | 1;100 |
| F4/80 | Bio-Rad | MCA497BB | Rat | 1;100 |

| Western blotting (WB) | | | | |
|---|---|---|---|---|
| pERK | Cell Signaling Technology | 4695S | Rabbit | 1;2000 |
| ERK | Cell Signaling Technology | 4370S | Goat | 1;1000 |
| pAkt | Cell Signaling Technology | 4060S | Rabbit | 1;2000 |
| Akt | Cell Signaling Technology | 4685S | Rabbit | 1;1000 |
| pSTAT1 | Cell Signaling Technology | 4649S | Rabbit | 1;1000 |
| STAT1 | Cell Signaling Technology | 14994S | Rabbit | 1;1000 |
| pSTAT3 | Cell Signaling Technology | 9145S | Rabbit | 1;1000 |
| STAT3 | Cell Signaling Technology | 4904S | Rabbit | 1;1000 |
| human IFNAR1 | Abcam | ab124764 | Rabbit | 1;1000 |
| mouse IFNAR1 | Abcam | ab97701 | Rabbit | 1;500 |
| pNF-κB p65 | Cell Signaling Technology | 3033 | Rabbit | 1;100 |
| NF-κB p65 | Cell Signaling Technology | 8242 | Rabbit | 1;100 |
| β-Actin | Sigma-Aldrich | A5441 | mice | 1;5000 |

Primers for qPCR

| Gene | Host | F | R |
|---|---|---|---|
| IFNA1 | human | GCCTCGCCCTTTGCTTTACT | CTGTGGGTCTCAGGGAGATCA |
| IFNA13 | human | CTATGATGGCCTCGCCCTTT | CTGGAGCCTTCTGGAACTGG |
| IFIT2 | human | GCAAGCTACCGTCTGGACA | CTTGCCTCAGAGGGTCAATG |
| IFIT3 | human | GAACATGCTGACCAAGCAGA | CAGTTGTGTCCACCCTTCCT |
| TRIM22 | human | AGAAGCTGGAAGATGACATCA | AGCTGCTGCCAGGTTATC |
| OASL | human | CTGATGCAGGAACTGTATAGCAC | CACAGCGTCTAGCACCTCTT |
| DDX60 | human | CAGCTCCAATGAAATGGTGCC | CTCAGGGGTTTATGAGAATGCC |
| TRAIL | human | TGCGTGCTGATCGTGATCTTC | GCTCGTTGGTAAAGTACACGTA |
| IFNG | mice | GCCACGGCACAGTCATTGA | TGCTGATGGCCTGATTGTCTT |

### Supplemental Table S2: Neutralizing antibody and plasmid information

| Antibody | Company | Catalog # | Clone |
| --- | --- | --- | --- |
| Anti-human IFNAR | Millipore | MAB1155 | MMHAR-2 |
| Anti-mouse CTLA4 | Bio X Cell | BE0164 | 9D9 |
| Anti-mouse PD1 | Bio X Cell | BP0146 | RMP1-14 |
| Anti-mouse IFNAR1 | Bio X Cell | BE0241 | MAR1-5A3 |

| sh-construct | Company | Catalog # |
| --- | --- | --- |
| Non-Target shRNA Control Plasmid | Sigma-Aldrich | #SHC016 |
| human IFNAR1 shRNA Plasmid | Sigma-Aldrich | TRCN0000059013 |
| | Sigma-Aldrich | TRCN0000059014 |
| | Sigma-Aldrich | TRCN0000428623 |
| mouse IFNAR1 shRNA Plasmid | Sigma-Aldrich | TRCN0000374694 |
| | Sigma-Aldrich | TRCN0000301483 |
| human STAT1 shRNA Plasmid | Sigma-Aldrich | TRCN0000280021 |
| | Sigma-Aldrich | TRCN0000004267 |
| EF.STAT3C.Ubc.GFP | Addgene | #24983 |
| pLVE-eGFP | Addgene | #52581 |
